# State-dependent structured sparsification of cortical signal-transmission networks enhances visual coding

**DOI:** 10.64898/2026.07.19.737501

**Authors:** Jiaji Zhu, Xiaoxuan Jia

## Abstract

The brain can rapidly adjust sensory processing according to behavioral context from moment to moment without altering its underlying anatomical wiring. Such flexibility is thought to arise from dynamic reconfiguration of the effective network through which signals propagate, yet the principles governing such reconfiguration and its consequences for coding remain unclear. Here, we exploited the distinct stationary and locomotion states within the same recording sessions and addressed this question by inferring directed, millisecond-scale signal-transmission networks at single-neuron resolution in behaving mice. Locomotion was accompanied by a counterintuitive structured sparsification of the inferred network: interactions became fewer but more temporally precise, while the remaining interactions were more local, modular, feature-specific, and feedforward. We then used theoretical analysis and controlled perturbations of multi-area rate models to systematically determine how each empirically observed component of this reorganization affects population coding. We found that sparsification reduced shared variability; local organization reduced signal–noise alignment; feature-specific interactions sharpened selectivity; and a more feedforward architecture accelerated decoding. These results provide an experimentally grounded mechanism by which behavioral state reorganizes neuronal interactions given the same sensory inputs, and suggest structured sparsification as an underlying principle of network reconfiguration that supports accurate and faster sensory coding.

## Introduction

The relationship between network architecture and neuronal population coding is a central question in systems and computational neuroscience. Anatomical connectivity constrains the possible routes through which spiking activity can propagate, but it does not uniquely determine the effective interactions engaged during sensory processing. Sensory and behavioral context can recruit different patterns of functional communication within the same anatomical substrate (Tang et al., 2024; Bosman et al., 2012; Srinath et al., 2021). This flexibility provides an opportunity to study the network–coding relationship in vivo: coordinated changes in effective interactions and coding performance within the same circuit can reveal which aspects of network organization are functionally consequential. However, which network features are reconfigured in behaving animals, and how their reorganization shapes coding fidelity and speed, remain unclear.

Theoretical and computational work has identified candidate circuit properties through which such reorganization could affect coding and population dynamics. Recurrent circuit models show that connectivity can organize shared trial-to-trial variability (Rosenbaum and Doiron, 2014; Rosenbaum et al., 2017; Huang et al., 2019, 2022), which limits Fisher information when aligned with stimulus-sensitive population directions (Moreno-Bote et al., 2014; Kanitscheider et al., 2015b). Low-rank connectivity links network topology to low-dimensional dynamics and computational capacity (Mastrogiuseppe and Ostojic, 2018), whereas local recurrent connectivity and connectivity motifs regulate activity dimensionality (Recanatesi et al., 2019; Dahmen et al., 2023; Shao and Ostojic, 2023; Shao et al., 2025). Clustered architectures organize ensemble dynamics and trial-to-trial variability (Litwin-Kumar and Doiron, 2012; Mazzucato et al., 2015), while state-dependent changes in intrinsic gain or metastable dynamics can alter stimulus discriminability and processing speed (Mazzucato et al., 2016, 2019; Wyrick and Mazzucato, 2021; Papadopoulos et al., 2026). Other related work shows that transient synchronization can flexibly route activity through fixed connectivity (Palmigiano et al., 2017). Together, these studies establish that circuit properties can influence population coding through shared variability, representational geometry, and temporal dynamics. Much of this work begins with a candidate property or mechanism and asks what computational consequences it produces. The complementary challenge is to identify the architecture of network reorganization expressed in vivo in behaving animals and use those observed network properties to define mechanistic tests of their contributions to coding fidelity and speed.

Naturally alternating stationary and locomotion states provide a tractable setting to address this challenge. Because these conditions can occur within the same recording sessions with the same visual inputs, changes in effective interactions and coding can be compared between states while anatomical connectivity remains relatively fixed. Locomotion modulates visually evoked responses and population correlation structure, and can improve the fidelity and speed of visual coding (Niell and Stryker, 2010; Ayaz et al., 2013; Vinck et al., 2015; Dadarlat and Stryker, 2017; Clancy et al., 2019; Christensen and Pillow, 2022; Wyrick and Mazzucato, 2021; Horrocks et al., 2024). It also alters pairwise neuronal correlations, the coupling of individual neurons to distributed cortical activity, and microscale functional circuitry (Dadarlat and Stryker, 2017; Clancy et al., 2019; Horrocks et al., 2024). We therefore used the stationary vs locomotion contrast not simply to characterize a behavioral-state effect on network reorganization, but as an empirical setting to study the network–coding relationship. Specifically, we asked whether changes in visual coding are accompanied by a coordinated reorganization of signal propagation networks, what form this reconfiguration takes, and mechanistically how its empirically observed components influence coding fidelity and speed.

Using simultaneous Neuropixels recordings from six visual cortical areas in the Allen Brain Observatory (Siegle et al., 2021), we inferred directed, millisecond-scale neuronal interactions from short-latency, jitter-corrected spike-train cross-correlograms and identified sharp peaks with brief latency. This approach emphasizes fast-timescale interactions while reducing contributions from stimulus locking and slower co-modulation (Smith and Kohn, 2008; Jia et al., 2022). The resulting graphs were referred as fast-timescale signal-transmission networks. We found that improved coding during locomotion was associated with structured sparsification of these networks. Overall interaction density decreased, but the remaining interactions were more temporally precise, local, feature-specific, and feedforward-biased, formed a more modular network, and were redistributed toward weakly selective neurons. Guided by these empirically observed dimensions of reorganization, we systematically manipulated each network feature in multi-area rate models and quantitatively evaluated its influence on coding using linear Fisher information analysis and population dynamics. The model perturbations revealed distinct computational contributions: sparsification reduced shared variability; local and modular organization reduced signal–noise alignment; feature-specific interactions and increased input to weakly selective neurons sharpened stimulus selectivity; and greater feedforward bias straightened population trajectories and accelerated decoding. Together, these results show that the same anatomical circuit can express distinct fast interaction networks across behavioral conditions during repeated presentation of the same visual stimuli. They identify structured sparsification as an empirically grounded network principle through which fewer, better-organized interactions can support more accurate and faster sensory coding.

## Results

Consider a neural population with signal-transmission matrix *W*, driven by sensory inputs *I* (*θ;*) and internal noise *ξ* (Stringer et al., 2019; Musall et al., 2019), where *θ* indicates the stimulus parameter. The structure of *W* shapes the population responses and therefore modulates information-coding capacity and dynamics. How locomotion reconfigures *W* and how this network reconfiguration modulates the coding capacity and speed are the central questions we address in this study (Fig. 1A).

**Fig. 1.**
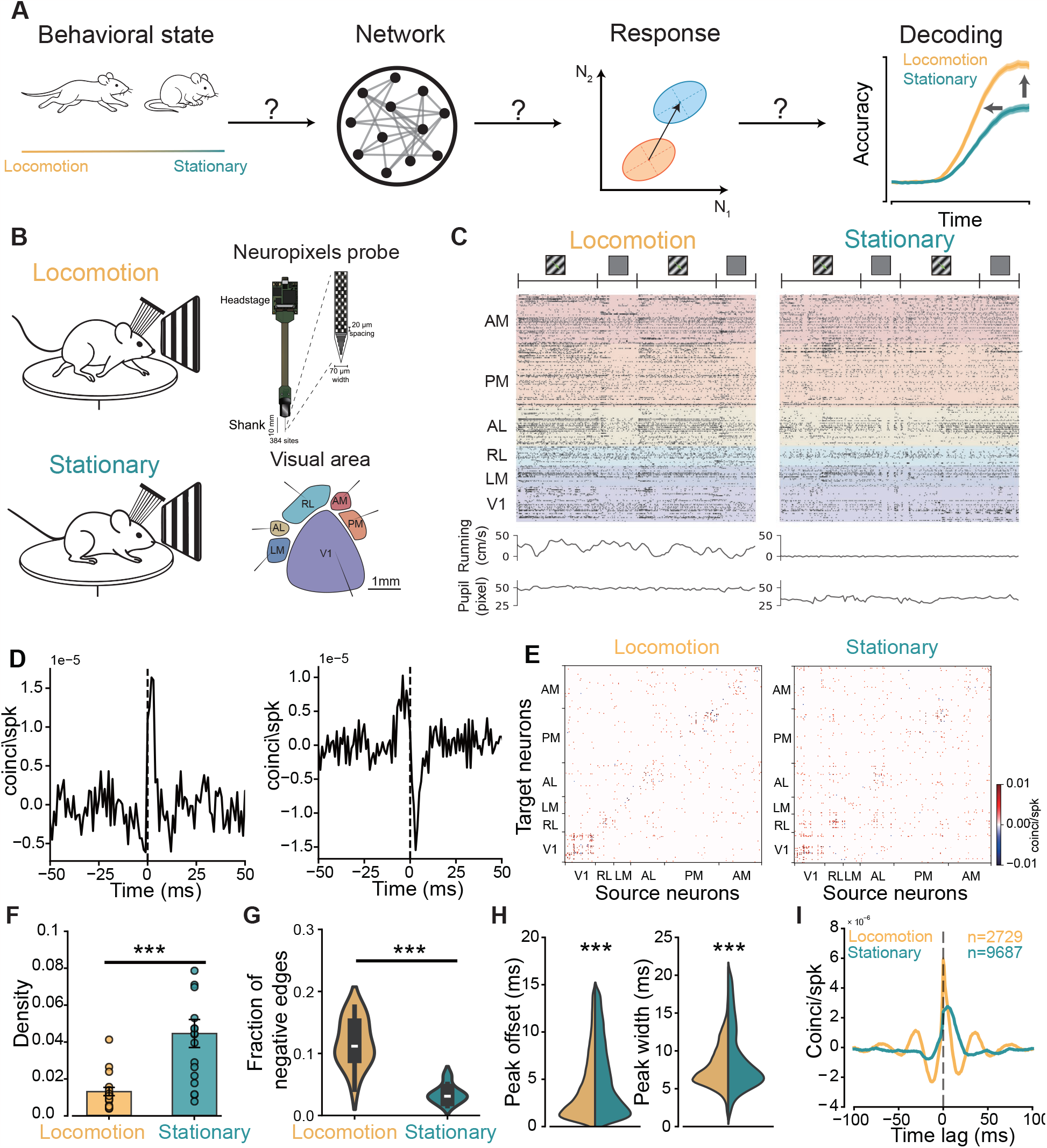
Locomotion reduces the network density. **A**, Schematic illustration of the key questions addressed in this study. We investigated how behavioral state altered network organization in the visual cortex, how network reconfiguration reshaped neural responses, and how changes in neural responses affected information decoding. **B**, Schematic of data collection using Neuropixels probes inserted across six visual cortical areas (AM, PM, AL, RL, LM, and V1). **C**, Example raster plot of simultaneously recorded units from six areas during drifting grating stimulation, shown for both locomotion and stationary states, with the running-speed and pupil size trace below. **D**, Example positive CCG trace (left) and negative CCG trace (right). **E**, Example adjacency matrix during locomotion (left) and stationary states (right). **F**, Network density during locomotion and stationary states in response to drifting grating stimuli. Error bars indicate SEM; points indicate individual mice (Wilcoxon signed-rank test, *n* = 17 mice, *p* = 1.52 × 10^−5^). Density was defined as the fraction of significant edges relative to all possible edges. **G**, Fraction of negative edges during locomotion and stationary states in response to drifting grating stimuli. (Wilcoxon signed-rank test, *n* = 17 mice, *p* = 1.52 × 10^−5^). Fraction of negative edges was defined as the fraction of significant negative edges among all significant edges. Violin plots show the distribution; white lines indicate medians and dark boxes the interquartile range. **H**, CCG peak offset (Wilcoxon signed-rank test, *n* = 17 mice, *p* = 1.52 × 10^−5^) and peak width (Wilcoxon signed-rank test, *n* = 17 mice, *p* = 0.0006). **I**, Average CCG trace for significant pairs during drifting gratings stimuli.

We analyzed the Allen Brain Observatory Neuropixels dataset (Siegle et al., 2021), which contains simultaneous recordings of spiking activity from six cortical visual areas (Fig. 1B; V1, LM, RL, AL, PM, AM) and two thalamic nuclei (LGN, LP) in mice passively viewing a variety of artificial and natural visual stimuli, while monitoring running speed and pupil size (Fig. 1B, C). We classified trials as locomotion or stationary based on mean running speed (threshold: 1 cm/s) to determine different behavioral states. In each behavioral state, we inferred pairwise connections using jitter-corrected cross-correlograms (CCGs) (Jia et al., 2013; Siegle et al., 2021; Smith and Kohn, 2008). Connections were inferred between neuron pairs whose jitter-corrected CCGs exhibited narrow, asymmetric peaks at short, biologically plausible lags: positive peaks reflect increased firing probability of the target neuron after the source neuron fires, whereas negative peaks reflect decreased firing probability (Fig. 1D). Since these statistical couplings have been suggested to reflect signal transmission efficacy (Smith and Kohn, 2008; Kohn et al., 2016; Jia et al., 2013; Siegle et al., 2021), potentially including monosynaptic, polysynaptic and common-input-mediated interactions, we refer to the resulting networks as signal-transmission networks (Fig. 1E). Unlike spike-count correlation or noise-correlation networks, these graphs emphasize asymmetric, short-latency spike-train dependencies.

### Locomotion reorganizes the signal-transmission network through structured sparsification

Locomotion has been shown to reduce correlated variability in the visual cortex (Dadarlat and Stryker, 2017). Because correlated variability and functional coupling are both shaped in part by circuit architecture (Kohn et al., 2016), this raised the question of whether locomotion influences the signal-transmission network similarly. To address this, we quantified network density *d* as the fraction of inferred connections among all possible directed connections, 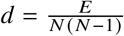, where *E* denotes the number of inferred connections and *N* is the number of neurons. We found that the density was consistently lower during locomotion than during stationary periods for drifting gratings (Fig. 1F, Wilcoxon signed-rank test, *p* = 1.52× 10^−5^), static gratings, natural scenes, and natural movies (Extended Data Fig. 1B). This decrease was robust across visual areas (Fig. 2C). These results indicated that locomotion sparsified the signal-transmission network, consistent with previous reports that locomotion decorrelates neural activity (Dadarlat and Stryker, 2017; Erisken et al., 2014; Horrocks et al., 2024). Our findings further extended these observations beyond V1 to multiple visual areas and moved beyond noise correlations to directed, spike-timing-based signal transmission. Notably, the sparsification was sign-biased: the reduction in density was restricted to positive connections (Extended Data Fig. 2A); Negative-connection density was unchanged for drifting and static gratings, and was even higher during locomotion for natural scenes and natural movies (Extended Data Fig. 2B). Therefore the fraction of negative connections was higher during locomotion (Fig. 1G, Wilcoxon signed-rank test, *p* = 1.52 × 10^−5^). Beyond this network sparsification, locomotion accelerated and sharpened signal transmission, as reflected by shorter offsets and narrower widths of CCG peaks (Fig. 1H, I; peak offset: Wilcoxon signed-rank test, *n* = 17 mice, *p* = 1.52 × 10^−5^; peak width: Wilcoxon signed-rank test, *n* = 17 mice, *p* = 6.0 × 10^−4^; Extended Data Fig. 3A, B).

**Fig. 2.**
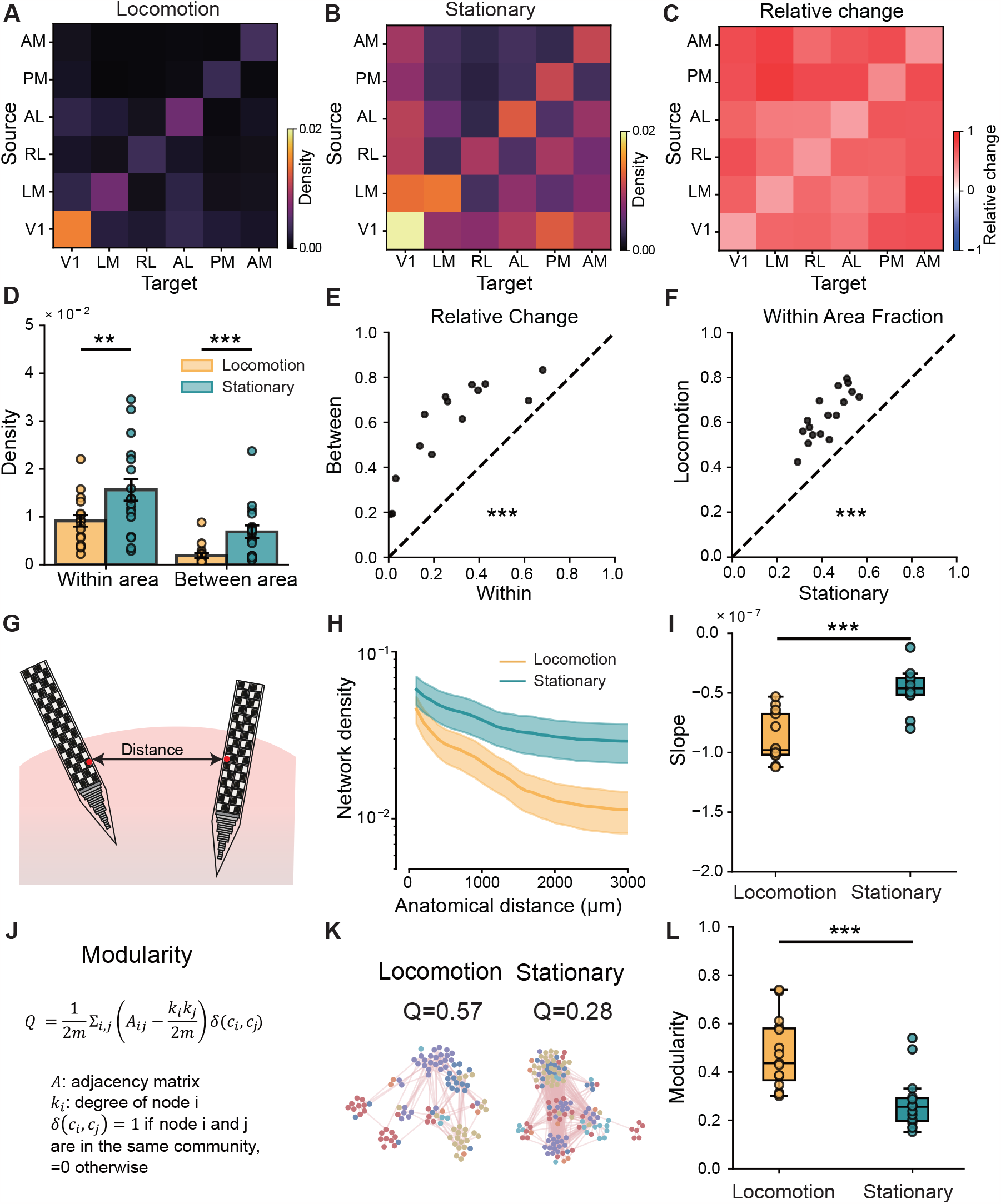
Locomotion localizes connectivity and increases network modularity. **A**, Network density of connections among six visual areas during locomotion. **B**, Same as in **A**, but during the stationary state. **C**, Relative change in network density across the six visual areas. Relative change in density was defined as (*d*_*stationary*_ −*d*_*locomotion*)_ / (*d*_*stationary*_ +*d*_*locomotion*_), where *d*_*stationary*_ and *d*_*locomotion*_ denote network density during stationary and locomotion states. **D**, Network density of within-area and between-area connections during drifting grating stimuli. Error bars indicate SEM, and points indicate individual mice (Wilcoxon signed-rank test, *n* = 17 mice, within-area: *p* = 1.52 ×10^−3^; between-area: *p* = 1.52 ×10^−5^). **E**, Relative changes in within-area density and between-area density. Points represent individual mice (paired *t*-test, *n* = 17 mice, *p* = 1.81 ×10^−10^). **F**, Within-area fraction of connections, defined as the number of connections between neurons in the same area divided by the total number of connections (paired *t*-test, *n* = 17 mice, *p* = 2.46 ×10^−10^). **G**, Schematic illustration of anatomical distance between neurons. **H**, Network density as a function of anatomical distance. Shading denotes ± SEM. (*n* = 17 mice). **I**, Slopes of curves in **H** for each mouse. Box plots show the median (center line), interquartile range (box), and whiskers extending to 1.5 IQR; points represent mice (paired t-test, *n* = 17 mice, *p* = 1.43 ×10^−8^). **J**, Schematic illustration of network modularity. **K**, Example network visualization from a representative mouse during locomotion (left) and stationary (right) states. **L**, Network modularity across conditions. Box plots show the median (center line), interquartile range (box), and whiskers extending to 1.5× IQR; points represent mice (Wilcoxon signed-rank test, *n* = 17 mice, *p* = 6.10 × 10^−5^).

Naive cross-correlograms are sensitive to firing rate (Stark and Abeles, 2009), which is strongly modulated by behavioral state (Niell and Stryker, 2010). However, our jitter-corrected CCGs were normalized by the geometric mean of the two neurons’ firing rates and further corrected by jittering (Methods) to minimize the influence of firing rates fluctuations beyond the jitter window. To confirm that the density decrease was not driven by firing rate modulation, we performed a rate-matching control in which we randomly subsampled spikes to equate firing rates across states while preserving spike-timing structure (Extended Data Fig. 4A). Networks constructed from rate-matched spike trains still showed reduced density during locomotion (Extended Data Fig. 4C), indicating that this effect cannot be explained solely by firing-rate changes.

Taken together, locomotion systematically sparsified the signal-transmission network *W*, preferentially eliminating positive over negative connections. This pattern suggested that the sparsification was structured rather than random. In the following sections, we characterize which features of *W* undergo structured sparsification and reveal their effects on information coding and dynamics.

### Locomotion shifts the network towards local and modular organization

Anatomical organization constrains neuronal connectivity, such that the connection probability between pairs of neurons is generally higher when the neurons are in the same area and decreases with anatomical distance (Markov et al., 2013, 2014). This spatial constraint shapes noise covariance (Rosenbaum et al., 2017; Huang et al., 2019), determines the stability of neural dynamics (Rosenbaum and Doiron, 2014; Huang et al., 2019), modulates the intrinsic timescale (Zeraati et al., 2023), and influences information-coding capacity (Huang et al., 2019). We asked whether locomotion-induced sparsification was spatially structured by testing whether its magnitude differed between within-area and between-area connections and varied with anatomical distance. To quantify the magnitude of sparsification, we defined a relative change of density as (*d*_*stationary*_ − *d*_*locomotion*_) / (*d*_*stationary*_ +*d*_*locomotion*)_, where *d*_*locomotion*_ and *d*_*stationary*_ denote density during locomotion and stationary states, respectively. We found that locomotion sparsified connection across all six visual areas, yet between area connections were sparsified more drastically (Fig. 2A–C): the relative change was significantly higher for between-area than for within-area connections (Fig. 2D, E, Extended Data Fig. 5A, B) and the fraction of within-area connections increased during locomotion (Fig. 2F, paired t-test, *p* = 2.46 × 10^−10^). Similarly, during locomotion, density declined more steeply with distance between recorded units (Fig. 2G-I, Extended Data Fig. 5C, D). These results revealed the spatial structure of locomotion-induced sparsification: it preferentially removed long-range connections, shifting the signal-transmission network toward a more localized organization.

We examined whether this localization was accompanied by higher network modularity (Deco et al., 2015; Cohen and D’Esposito, 2016; Sporns, 2013; Baum et al., 2017). We quantified modularity using the Louvain community-detection algorithm (Blondel et al., 2008), which captures the extent to which a network is partitioned into modules with stronger within-module than between-module connectivity (Fig. 2J, K). The resolution parameter of the detection method was chosen to maximize the deviation from a degree- and density-matched null model (Methods; Extended Data Fig. 7A). Modularity was significantly higher during locomotion (Fig. 2L, Wilcoxon signed-rank test, *p* = 6.10 × 10^−5^), and this increase was not solely due to differences in density. A null-model-corrected modularity score, obtained by subtracting the modularity of a pair-preserving null model, remained significantly higher during locomotion (Extended Data Fig. 7C).

Similar structured sparsification was observed across different significant-CCG-peak thresholds (Extended Data Fig. 6A–C). These results indicate that sparsification was spatially structured, biasing the network towards a more localized organization. Between-area and long-range connections were preferentially eliminated while short-range and within-area connections were comparatively preserved, shifting the network from a globally integrated architecture towards a more localized and modular organization.

### Localized networks improve coding by reducing information-limiting correlations

To test whether and how sign-biased and spatially localized sparsification can improve population coding, we built a two-layer rate-based network with independently tunable connection density, spatial range, and edge sign fraction (Fig. 3A–C). Each layer contained 200 neurons, with a 4:1 ratio of excitatory to inhibitory units. To approximate hierarchical visual processing (Siegle et al., 2021), we used a two-layer hierarchical model in which stimuli entered the lower layer and propagated to the higher layer, with spatially structured connectivity defined by distance-dependent connection probabilities. We computed linear Fisher information (Fig. 3D) to quantify the amount of stimulus information accessible to an optimal linear decoder (Moreno-Bote et al., 2014):

**Fig. 3.**
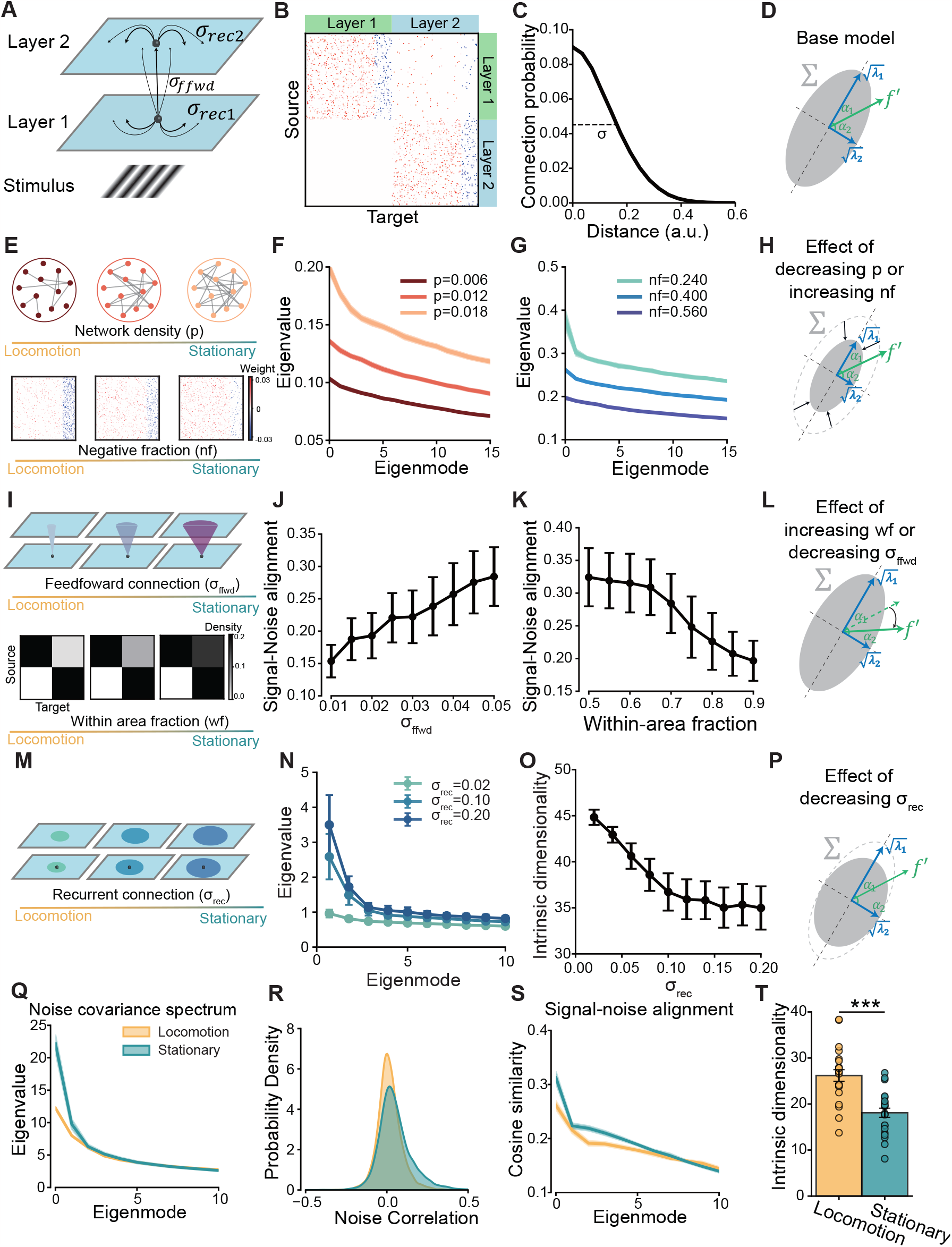
Localized networks reshape noise covariance to reduce information-limiting correlation. **A**, Schematic of a network model. Neurons in each layer are arranged in a 2D plane. The network consists of two layers. **B**, Example adjacency matrix of the network. Red and blue entries indicate positive and negative values, respectively. **C**, Distance dependence of connection probability. **D**, Fisher information was decomposed into the magnitude of the signal (∥ *f*′ ∥), the magnitude of noise covariance 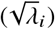, and the alignment between signal and dominant noise directions (*α*_1_). **E**, Top, increasing network density from sparse to dense. Bottom, decreasing the fraction of negative connections. **F**, Eigenvalue spectrum of the noise covariance matrix at different network densities. Shading denotes SEM across 20 independently generated network realizations and is too small to be visible. **G**, Eigenvalue spectrum of the noise covariance matrix at different fractions of negative connections (*n f*). Shading denotes SEM across 20 independently generated network realizations and is too small to be visible. **H**, Reduced density or a higher fraction of negative connections reduced noise covariance. **I**, Top, increasing the spatial scale of feedforward connections from narrow to broad. Bottom, decreasing the within-area fraction of connections. **J**, Effect of feedforward connection spatial scale on the alignment between signal direction ( *f*′) and dominant noise direction (the dominant eigenvector of the noise covariance matrix). Alignment is defined as the cosine similarity, which is higher when signal and noise are more similarly oriented. Error bars indicate SEM. (*n* = 20 repeats). **K**, Effect of within-area fraction on the alignment between signal direction ( *f*′) and dominant noise direction. Error bars indicate SEM. (*n* = 20 repeats). **L**, Within-area, localized networks reduced signal–noise alignment. **M**, Increasing the spatial scale of recurrent connections from narrow to broad. **N**, Eigenvalue spectrum of the noise covariance matrix at different spatial scales of recurrent connectivity. Error bars indicate SEM. (*n* = 20 repeats). **O**, Intrinsic dimensionality decreased as the spatial scale of recurrent connections increased. Error bars indicate SEM. (*n* = 20 repeats). **P**, Localized networks increased the intrinsic dimensionality of noise. **Q**, Eigenvalue spectrum of the noise covariance matrix in the experimental data. Shaded areas indicate SEM. (*n* = 17 mice). **R**, Distribution of noise correlations across states (*n* = 17 mice). **S**, Signal–noise alignment, quantified as the cosine similarity between *f*′ and covariance eigenvectors, in the experimental data. Shaded areas indicate SEM (*n* = 17 mice). **T**, Intrinsic dimensionality of neural responses. Points represent individual mice. Error bars indicate SEM. (Paired t-test, *p* = 8.86 × 10^−7^, *n* = 17 mice).

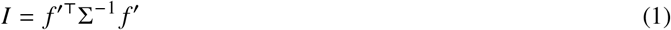

where Σ is the noise covariance matrix and *f*′ = *df* (*θ*) /*dθ* denotes the derivative of the mean population response *f* with respect to the stimulus parameter *θ*. Larger linear Fisher information corresponds to higher population decoding accuracy. Fisher information depends on three components: the magnitude of noise covariance, signal strength and signal-noise alignment (Methods). We examined how structured sparsification changes each of these components.

As previously reported (Dadarlat and Stryker, 2017), locomotion reduced the shared variability of neural activity. We asked whether sparsification and sign-biased reconfiguration of networks contribute to this effect. To test this, we varied the total density and measured the noise covariance (Fig. 3E). Decreasing density reduced the eigenvalues of the noise covariance matrix (Fig. 3F), suggesting that sparsification reduced shared variability. We further tested the effect of an increased negative fraction. At fixed density (*p* = 0.1), increasing the negative fraction also reduced eigenvalues (Fig. 3G). These results suggest that sparsification and an increased negative fraction reduce the magnitude of noise covariance, thereby increasing Fisher information (Fig. 3H).

Linear Fisher information depends not only on the magnitude of shared noise, but also on its alignment with the stimulus-related signal (Moreno-Bote et al., 2014). Noise becomes information-limiting when it lies along the signal direction, which we quantified as the cosine similarity between the signal derivative ( *f*′) and the dominant noise direction (*u*_1_). Feedforward projections are a source of information-limiting correlation, because signal and noise propagate through the same pathways and are therefore difficult for downstream populations to disentangle (Kanitscheider et al., 2015b). We hypothesized that more spatially restricted feedforward projections would reduce signal–noise alignment by reducing shared feedforward circuitry. We also hypothesized that increasing the within-area fraction of connections would weaken information-limiting correlation by reducing the relative contribution of feedforward input.

To test these predictions, we varied the spatial scale of feedforward connectivity σ_ffwd_ (Fig. 3I). Decreasing σ_ffwd_ reduced the alignment between signal and dominant shared noise (quantified by cosine similarity) (Fig. 3J), indicating weaker signal–noise alignment. Increasing the within-area fraction produced a similar effect (Fig. 3K). These observations were robust to a range of parameters (Extended Data Fig. 8A, B). Because locomotion shifts the network towards more within-area and spatially localized connectivity structure (Fig. 2), these results suggest that locomotion reduces the information-limiting effect of noise correlations (Fig. 3L). Consistently, alignment between *f*′ and the leading eigenvectors of the noise covariance was lower during locomotion in the experimental data (Fig. 3S).

Unlike the spatial scale of feedforward connections, the spatial scale of recurrent connections primarily shapes network integration (Cohen and D’Esposito, 2016; Deco et al., 2015; Rosenbaum et al., 2017). We hypothesized that broader recurrent connectivity would promote population-wide shared dynamics, concentrating variability into a few dominant modes and reducing the effective dimensionality of population activity. To test this, we varied the recurrent spatial scale σ_rec_ (Extended Data Fig. 8C, Fig. 3M). Narrower recurrent connectivity substantially reduced the first few dominant eigenvalues of the noise covariance, while leaving the smaller eigenvalues largely unchanged (Fig. 3N). As a result, intrinsic dimensionality increased (Fig. 3O).

Consistent with the simulations, noise correlation decreased during locomotion (Fig. 3R) and the leading eigenvalue decreased during locomotion in the mouse visual cortex (Fig. 3Q) and intrinsic dimensionality was higher during locomotion (Fig. 3T). Increased intrinsic dimensionality does not by itself imply higher Fisher information, because if *f*′ were orthogonal to the dominant noise modes, changes in their eigenvalues would have little effect on coding. In the mouse visual cortex, however, the dominant eigenmodes of the noise covariance were also most aligned with *f*′, as indicated by the highest cosine similarity (Fig. 3S). The increase in dimensionality (Fig. 3T) thus reflects selective reduction of noise along signal-relevant dimensions thereby also increasing Fisher information (Fig. 3P).

These results demonstrate that sign-biased and spatially ordered sparsification reshapes noise covariance to increase Fisher information through multiple mechanisms. Reduced density and an increased negative fraction suppress shared variability, reducing the magnitude of noise covariance. More within-area and more localized feedforward connectivity reduces signal–noise alignment. Narrower recurrent connectivity redistributes noise away from signal-relevant dimensions, reducing the information-limiting effect of noise correlations.

### Locomotion reorganizes the network to enhance feature selectivity

Linear Fisher information is proportional to signal magnitude ∥*f*′∥, which is large when neurons respond selectively (Moreno-Bote et al., 2014). Neurons with similar tuning are more likely to be connected (Cossell et al., 2015; Ding et al., 2025; Ko et al., 2011), and such feature-specific connectivity can enhance neuronal selectivity (Huang et al., 2022). We therefore asked whether sparsification was structured along tuning similarity. We computed tuning similarity as the difference in preferred direction (drifting gratings) or orientation (static gratings) and then quantified how density changed with tuning similarity. This density was normalized by the overall network density to account for differences in overall density across states, producing a relative density measurement. In both states, relative density generally decreased with increasing tuning difference (Fig. 4A, B), consistent with a feature-specific connectivity pattern. Neurons with identical tuning preferences exhibited higher relative density during locomotion for drifting gratings and static gratings (Fig. 4A, B). The same effect was observed when tuning similarity was quantified using signal correlations (Extended Data Fig. 9A, B). These results indicated that sparsification was structured along tuning similarity, producing a more feature-specific network during locomotion.

**Fig. 4.**
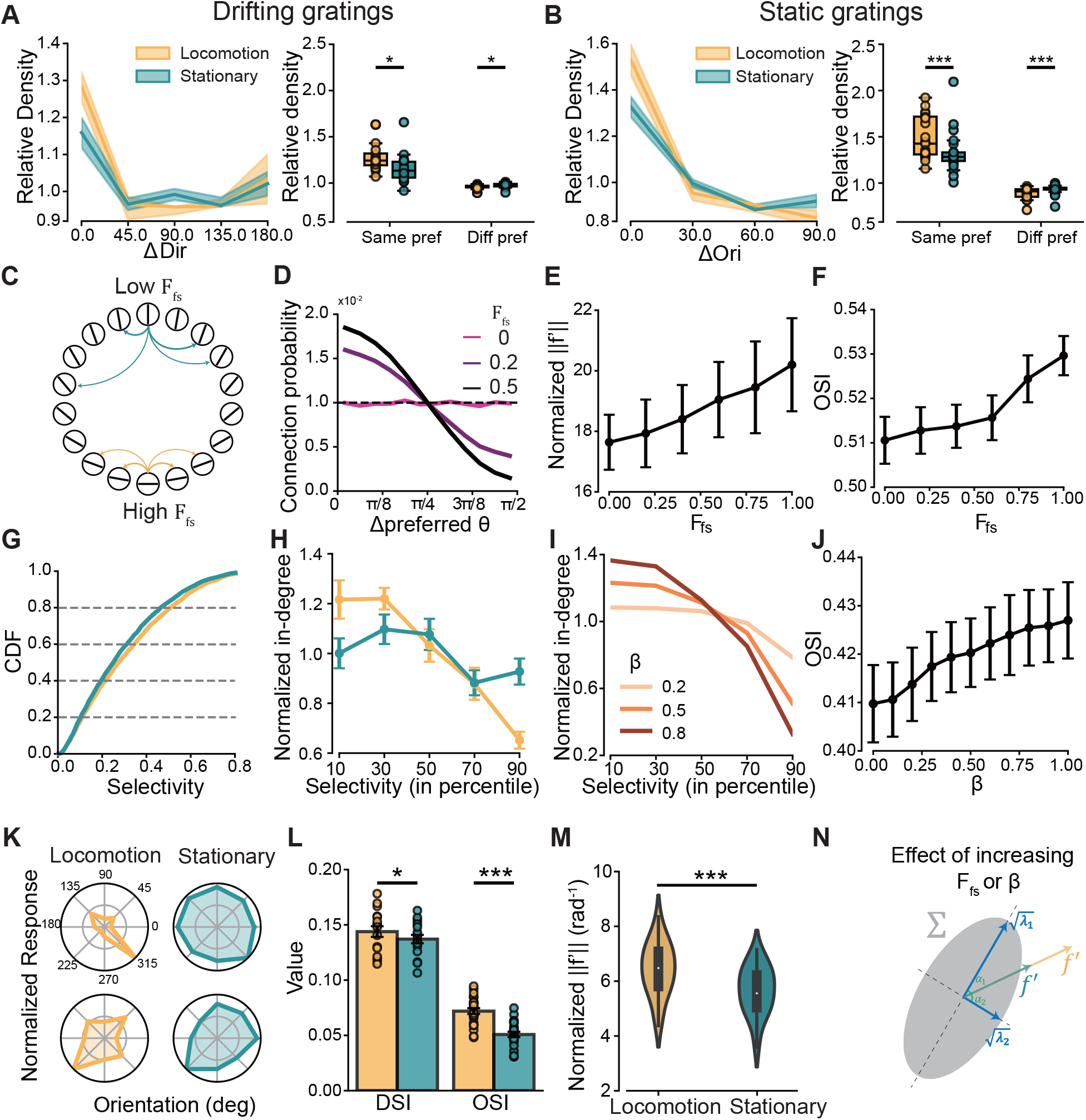
Locomotion increases feature selectivity through network reconfiguration. **A**, Left: Relative connection density between neuron pairs with different direction preferences during drifting grating stimulation. Shaded regions indicate mean ± SEM (*n* = 17 mice). Right: Relative density of connections between neurons with the same direction preference (Δ*Dir* = 0) versus different direction preferences (Δ*Dir* ≠ 0). (same pref; paired *t*-test, *n* = 17 mice, *p* = 0.017; diff pref; paired *t*-test, *n* = 17 mice, *p* = 0.02). **B**, Same as in **A**, but based on orientation preference instead of direction preference (*n* = 24 mice). (same pref; paired *t*-test, *n* = 24 mice, *p* = 6.3 ×10^−5^; diff pref; paired *t*-test, *n* = 24 mice, *p* = 5.6 ×10^−5^). **C**, Illustration of feature specificity. Each circle on the ring represents a neuron, and the bar inside each circle indicates its preferred orientation. **D**, Relationship between tuning similarity and connection probability under different levels of feature specificity. **E**, Normalized *f*′ as a function of feature specificity in simulations. Error bars indicate SEM (*n* = 20 repeats). **F**, In simulations, the orientation selectivity index (OSI) increased with feature specificity. Error bars indicate SEM (*n* = 20 repeats). **G**, Cumulative distribution functions of the selectivity index in stationary and locomotion states. **H**, Normalized in-degree of neurons with different selectivity levels (percentile). Error bars indicate SEM (*n* = 17 mice). **I**, Normalized in-degree of neurons with different selectivity levels under different values of *β*. Shaded areas indicate SEM (*n* = 20 repeats), which is too small to be visible. **J**, Relationship between *β* and OSI when *F*_*fs*_ = 0.2. **K**, Example tuning curves across states for two neurons in the experimental data. Responses were normalized to the maximum stimulus-evoked response. **L**, Direction selectivity index (DSI) during drifting grating stimulation (paired *t*-test, *n* = 17 mice, *p* = 0.029) and orientation selectivity index (OSI) during static grating stimulation (paired *t*-test, *n* = 24 mice, *p* = 4.4 ×10^−11^). **M**, Normalized magnitude of ∥*f* ∥′ in the experimental data during drifting gratings stimuli. Neural responses were normalized by *z*-scoring (paired *t*-test, *n* = 17 mice, *p* = 7.1 ×10^−7^). **N**, Higher feature specificity *F*_*fs*_ and larger *β* increased signal magnitude.

Previous work has shown that feature-specific connectivity can enhance selectivity when orientation preference is organized in a pinwheel map and internal noise is negligible (Huang et al., 2022). However, in mouse visual cortex, neither assumption holds: orientation preference follows a salt-and-pepper organization (Ohki and Reid, 2007; Ringach et al., 2016; Ohki et al., 2005), and non-visual fluctuations contribute substantially to response variability (Stringer et al., 2019; Musall et al., 2019). To test whether feature-specific connectivity still improves selectivity in this regime, we defined connection probability to decrease as tuning similarity decreases, with *F*_*fs*_ controlling the degree of feature specificity (Fig. 4C, D). Increasing *F*_*fs*_ systematically sharpened population tuning, increasing both the orientation selectivity index and the signal-derivative magnitude (∥*f*′∥) (Fig. 4E, F, Extended Data Fig. 10A). Responses were normalized before computing *f*′, ensuring that the increase in ∥*f*′∥ reflected sharper tuning rather than a trivial consequence of elevated firing rates.

Locomotion also increased the portion of inputs to low-selectivity neurons while decreased the portion of inputs to highly selective neurons. To quantify this effect, we used the normalized in-degree, defined as the in-degree of neurons divided by the mean of in-degree across the population. Neurons were divided into five subpopulations according to their selectivity (Fig. 4G), spanning the lowest selectivity quintile (0–20th percentile) to the highest (80–100th percentile). During locomotion, more selective neurons showed lower normalized in-degree (Fig. 4H), indicating that locomotion preferentially reduced input projections to selective neurons. To simulate this effect in the model, we introduced a parameter *β* that modulates the probability of input projections to selective neurons in our simulations. This parameter determines how fast in-degree decays with increasing selectivity. A high value of *β* indicated fast decay, corresponding to fewer input projections to selective neurons and more input projections to less selective neurons (Fig. 4I and Methods). We found that, when connections are feature specific (*F*_*fs*_ > 0), orientation selectivity increased monotonically with increasing *β* (Fig. 4J, Extended Data Fig. 10B).

This is because an input projection enhances a target neuron’s selectivity only when it originates from a presynaptic neuron that is more selective and shares similar orientation preference (Methods). Highly selective neurons have few such presynaptic source neurons, so most of their inputs reduce selectivity; decreasing their input number therefore increases selectivity. In contrast, low-selectivity neurons have many more-selective presynaptic source neurons, so increasing their input number raises the contribution of selectivity-enhancing inputs and thereby increases selectivity (Extended Data Fig. 11A–C).

Consistent with previous reports (Dadarlat and Stryker, 2017; Christensen and Pillow, 2022) and our simulations, locomotion increased single-neuron selectivity in the experimental data: both the direction selectivity index (DSI, drifting gratings) and orientation selectivity index (OSI, static gratings) were significantly higher during locomotion (Fig. 4K, L). This sharpening was also reflected in an increased magnitude of the normalized signal derivative ∥*f*′∥ (Fig. 4M, paired t-test, *p* = 7.1 × 10^−7^).

These results revealed that sparsification was structured along two axes: it preferentially removed connections between dissimilarly tuned neurons and removed input projections to selective neurons. Both forms of structured sparsification sharpened tuning, increased signal magnitude, and improved coding capacity (Fig. 4N).

### Locomotion promotes feedforward organization and accelerates decoding

Beyond enhancing information coding, locomotion also accelerates information transmission (Niell and Stryker, 2010). Fast visual processing has been linked to feedforward network architecture. A feedforward network matched human performance in a rapid visual discrimination task (VanRullen, 2007), whereas models with greater feedback processing better predicted neural responses when categorization was slowed by challenging images (Kar et al., 2019). We asked whether structured sparsification could reconfigure feedforward and feedback organization. We quantified connection directionality between pairs of visual cortical areas using a directionality score measuring the relative prevalence of source-to-target versus target-to-source connections (Methods). During both states, the directionality scores from anatomically lower areas to higher area were generally positive (Fig. 5A), suggesting a feedforward network architecture. While during locomotion, the directionality scores (Methods) for lower-to-higher area pairs became more positive (Fig. 5B, C, paired t-test, *p* = 0.002), indicating strengthened feedforward structure during locomotion. This result was robust across alternative directionality measures, including reachability and canonical correlation analysis (Extended Data Fig. 12A–D).

**Fig. 5.**
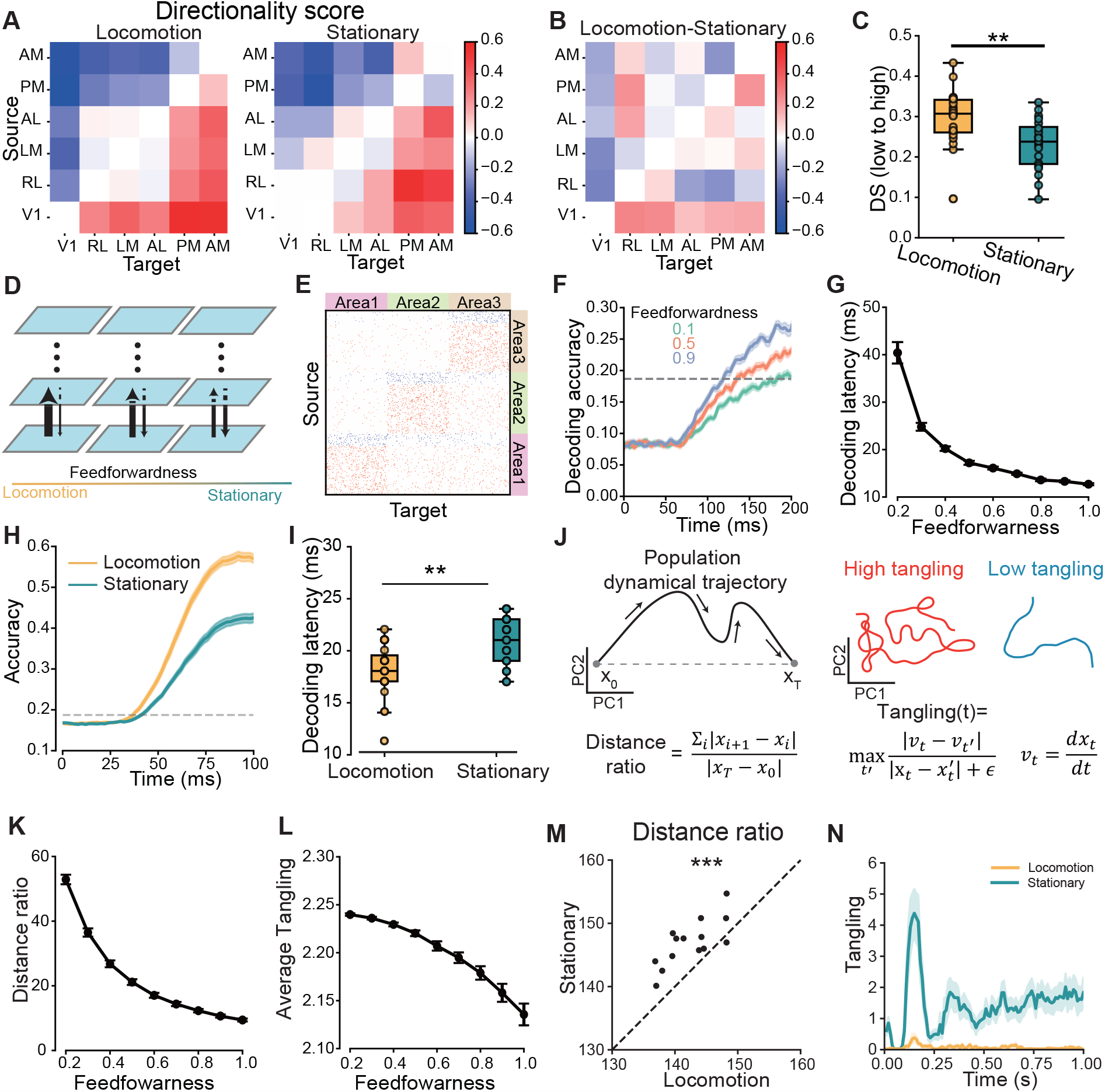
Feedforward reorganization supports faster visual decoding during locomotion. **A**, Directionality score calculated from the distribution of connection counts for each pair of cortical areas during locomotion (left) and stationary (right) states. Rows indicate source, columns indicate target. **B**, Difference in directionality score between the locomotion and stationary states. **C**, Directionality score (DS) for connections from anatomically lower to higher visual areas. The error distribution was estimated by bootstrap resampling over mice, with 20% of mice sampled in each repeat (paired *t*-test, *n* = 20 repeats, *p* = 0.002). **D**, Schematic of a multi-area network model. Feedforwardness gradually decreases from left to right. **E**, Example adjacency matrix at feedforwardness = 0.5. **F**, Time course of decoding accuracy in simulations under different values of feedforwardness (*n* = 20 repeats). Shading indicates SEM. The dashed line indicates the 95^th^ percentile of decoding accuracy under random guessing. **G**, Relationship between decoding latency and feedforwardness. Decoding latency was defined as the first time point at which decoding accuracy exceeded the 95th percentile of decoding accuracy under random guessing. (*n* = 20 repeats). **H**, Time course of decoding accuracy in the experimental data during static grating presentation. Shading indicates SEM (n=17 mice). **I**, Decoding latency in the experimental data. Box plots show the median (center line), interquartile range (box), and whiskers extending to 1.5 × IQR; points represent mice (Wilcoxon signed-rank test, *n* = 17 mice, *p* = 0.006). **J**, Schematic illustration of the distance ratio and tangling index. **K**, In simulations, the distance ratio decreased as feedforwardness increased. Error bars indicate SEM (*n* = 20 repeats). **L**, In simulations, the tangling index decreased as feedforwardness increased. Error bars indicate SEM (*n* = 20 repeats). **M**, Distance ratio in the experimental data during locomotion and stationary states (paired *t*-test, *n* = 17 mice, *p* = 3.5 × 10^−5^). **N**, Tangling index of neural trajectories in the experimental data during locomotion and stationary states. Shaded areas indicate SEM (*n* = 17 mice).

To test whether a more feedforward network can influence the speed of information coding, we constructed a three-layer hierarchical network model with both feedforward and feedback projections between areas (Fig. 5D, E). We defined feedforwardness as *d*_ff_/(*d*_ff_ + *d*_fb_), where *d*_ff_ and *d*_fb_ are the feedforward and feedback densities, respectively. Feedforwardness ranged from 0 to 1, with 0.5 indicating equal feedforward and feedback densities and 1 indicating exclusively feedforward connectivity. Total inter-areal connection density was held constant at 0.1 while feedforwardness was systematically varied.

We presented gratings spanning nine orientations to the model (40 repeats each; 200 ms duration) and decoded stimulus orientation from the modeled neural population activity. Decoding latency was defined as the earliest time at which accuracy exceeded the 95th percentile of a shuffled control (threshold=0.19). Increasing feedforwardness monotonically reduced decoding latency (Fig. 5F, G, Extended Data Fig. 13A). Consistent with simulations, we found that decoding latencies were shorter during locomotion in mouse visual cortex (Fig. 5H, I). While faster decoding during locomotion has previously been attributed to gain modulation (Wyrick and Mazzucato, 2021), our results identify a complementary network-level mechanism, namely, a shift toward more feedforward inter-areal organization.

Increasing feedforwardness also produced more direct and less tangled population dynamics, which are associated with smoother, more noise-robust latent dynamics (Russo et al., 2018; Horrocks et al., 2024). We quantified this using two metrics (Fig. 5J and Methods): the distance ratio and the tangling index (Russo et al., 2018; Horrocks et al., 2024). In the model, increasing feedforwardness reduced both metrics (Fig. 5K, L, Extended Data Fig. 13B, C). Experimentally, we observed reductions in both metrics during locomotion (Fig. 5M, N), consistent with the model’s predictions and prior work (Horrocks et al., 2024).

These findings suggest that structured sparsification preferentially removed feedback connections, shifting visual cortex toward a more feedforward configuration. This feedforward shift enables more efficient signal propagation to higher-order areas, producing earlier decodable stimulus representations and more direct population dynamics. Together, these effects provide a network-level explanation for the faster information processing observed during locomotion. This reconfiguration may help animals respond more rapidly to visual cues during movement, supporting navigation, predator avoidance, and other survival-critical behaviors.

## Discussion

We investigated how a cortical circuit with essentially stable anatomical wiring can alter sensory coding by reconfiguring fast neuronal interactions. During repeated presentation of the same visual stimuli, stationary and locomotion states were associated with distinct single-neuron-resolution signal-transmission networks. This reconfiguration took the counterintuitive form of structured sparsification (Fig. 6A, B): inferred interactions became fewer overall but more temporally precise, and those that remained were more local, modular, feature-specific, and feedforward. Thus, locomotion was not associated with a uniform strengthening or weakening of cortical interactions, but with selective reorganization. Guided by these empirically observed dimensions of network changes, we manipulated each feature in multi-area rate models (Fig. 6C). The models revealed distinct computational contributions: sparsification and a larger fraction of negative interactions reduced shared variability; local and modular organization reduced signal–noise alignment; feature-specific interactions and redistributed input toward weakly selective neurons sharpened selectivity; and greater feedforward bias straightened population trajectories and accelerated decoding. These results link dynamic reorganization of fast neuronal interactions to both the fidelity and speed of population coding.

**Fig. 6.**
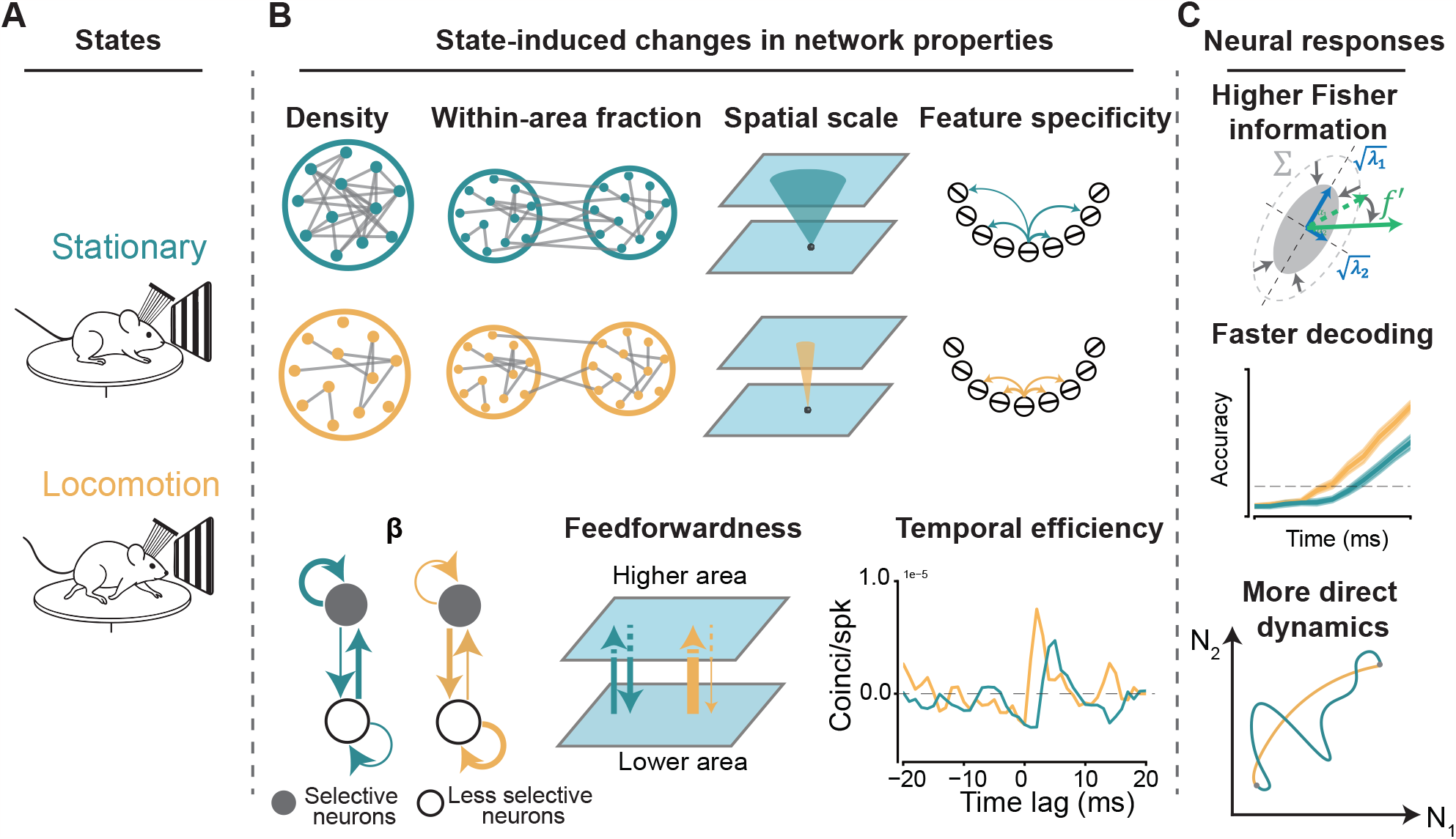
Overview of locomotion-induced reconfiguration of signal-transmission networks and its effects on Fisher information and neural dynamics. **A**, Illustration of stationary and locomotion states. **B**, Behavioral state-induced changes in network properties, including decreased density, a higher within-area fraction, a narrower spatial scale, heightened feature specificity, reduced incoming projection to highly selective neurons, increased feedforwardness, and improved temporal efficiency of signal transmission during the locomotion state. **C**, Effects of network changes on coding. Top: changes in the network properties shown in **B** result in reduced noise covariance, larger signal magnitude, and signal–noise misalignment, leading to increased Fisher information. Middle: network changes lead to higher decoding accuracy and shorter decoding latency during locomotion. Bottom: network reconfiguration straightened neural dynamics.

One important observation of this study is that improved coding was accompanied by a counterintuitive and coordinated change in network organization. There was no *a priori* reason to expect a sparser inferred network to support enhanced activity and coding accuracy. Behavioral and cognitive modulation can strengthen interareal synchronization or coupling (Gregoriou et al., 2009; Bosman et al., 2012; Ruff and Cohen, 2016), reduce within-area noise correlations (Mitchell et al., 2009; Cohen and Maunsell, 2009; Dadarlat and Stryker, 2017), or produce heterogeneous increases and decreases across neuronal pairs and cortical regions (Ruff and Cohen, 2014; Clancy et al., 2019). Enhanced responses or coding could therefore accompany stronger, weaker, or redistributed effective interactions without a coherent change in network architecture. The notable result was not sparsity alone, but structured sparsification: interactions became fewer overall, while those expressed during locomotion were more temporally precise and collectively formed a more local, modular, feature-specific, and feedforward-biased network. This coordinated pattern cannot be reduced to a uniform strengthening or weakening of cortical coupling and instead suggests that improved coding is associated with the selective organization of interactions into fewer, better-configured pathways.

The structured sparsification observed during locomotion (Fig. 1, Fig. 2, Fig. 4, and Fig. 5) may reflect a broader network mechanism for improving cortical information processing. Enhanced coding is also associated with attention (Cohen and Maunsell, 2009; Harris and Thiele, 2011; Saproo and Serences, 2010), arousal (Vinck et al., 2015; Papadopoulos et al., 2026), learning (Yan et al., 2014; Jeanne et al., 2013; Sanayei et al., 2018), and development (Smith et al., 2015; Chang and Fitzpatrick, 2022). Several of these conditions are accompanied by reduced correlated variability (Mitchell et al., 2009; Cohen and Maunsell, 2009; Gu et al., 2011; Jeanne et al., 2013; Sanayei et al., 2018; Smith et al., 2015; Chang and Fitzpatrick, 2022), but decorrelation alone does not imply structured sparsification. Our results make a more specific prediction: improved coding should be associated with fewer effective interactions together with greater temporal precision and selective organization of the interactions that are expressed. Testing this coordinated network signature across behavioral, cognitive, and developmental conditions will determine whether structured sparsification is a general mode of cortical computation or a reconfiguration specific to locomotion.

Beyond its possible generality, structured sparsification highlights the need to distinguish cortical organization at multiple levels. Anatomical connectivity defines a relatively stable set of possible pathways, whereas effective interactions reflect which routes are engaged in a particular sensory, cognitive, or behavioral context (Tang et al., 2024; Bosman et al., 2012; Srinath et al., 2021; Palmigiano et al., 2017). At the population level, interareal communication can be confined to low-dimensional communication subspaces, with feedforward and feedback interactions engaging distinct activity patterns (Semedo et al., 2019, 2022). How these population-level channels emerge from neuron-level, fast timescale signal propagation remains unresolved. Our study addresses an intermediate link in this multi-area network by characterizing state-dependent, fast interaction networks among individual neurons and using models to relate their observed reorganization to the geometry and temporal dynamics of population activity. A complete account will require determining how anatomical connectivity and state-dependent modulation generate these effective networks, establishing their causal influence in vivo, and testing whether the resulting information is accessible to downstream circuits and behavior.

A complementary feature of our approach is that the model perturbations were anchored to network changes observed in vivo. Previous theory has shown how prespecified variations in connectivity can regulate shared variability, activity dimensionality, population dynamics, and information (Rosenbaum and Doiron, 2014; Rosenbaum et al., 2017; Recanatesi et al., 2019; Huang et al., 2019, 2022). Here, the observed changes in density, locality, modularity, feature specificity, input distribution, and feedforward bias defined the axes of controlled perturbation. Their separate manipulation revealed a division of computational roles: density regulated the magnitude of shared variability; locality and modularity regulated signal–noise alignment; feature specificity and input redistribution regulated neuronal selectivity; and feedforward bias regulated trajectory geometry and decoding speed. These perturbations establish what each feature can do under the model assumptions, rather than demonstrating that the features are biologically independent or uniquely mediate the locomotion effect. Our localization result also differed from that obtained in a spatially ordered recurrent model (Huang et al., 2022). This difference may reflect the salt-and-pepper tuning and additional neuron-specific variability incorporated here (Ohki et al., 2005; Ohki and Reid, 2007; Ringach et al., 2016; Stringer et al., 2019; Musall et al., 2019), and emphasizes that the coding consequences of network structure depend on the joint organization of tuning, recurrent interactions, and variability.

The locomotion-associated increase in feedforward bias provides a complementary network-level account of faster sensory coding. Using the same Allen Brain Observatory Neuropixels dataset, Wyrick and Mazzucato found that locomotion was associated with reduced intrinsic gain during ongoing activity and earlier visual decoding; in a clustered recurrent model, reducing gain accelerated transitions into stimulus-selective metastable states (Wyrick and Mazzucato, 2021). Horrocks and colleagues further showed that locomotion was associated with more direct, less tangled population trajectories and faster stabilization of visual representations (Horrocks et al., 2024). Our models link these dynamical changes to effective network organization: increasing feedforward bias straightened population trajectories and shortened decoding latency. Gain modulation and network reorganization may therefore be coupled rather than competing mechanisms. Reduced gain could attenuate recurrent amplification and increase the relative influence of feedforward drive, whereas changes in recurrent or interareal interactions could alter effective gain and circuit intrinsic timescales. Because we did not manipulate these factors independently, we cannot determine their causal ordering. Our results instead identify greater feedforward bias as an observed effective-network change that is sufficient, under the model assumptions, to accelerate coding. Whether this reorganization also preferentially channels activity into population dimensions that effectively drive downstream areas remains to be tested.

The present study has several limitations. First, the inferred networks should be viewed as signal-transmission networks rather than anatomical networks or effective networks. Jitter-corrected cross-correlograms isolate fast-timescale spike-timing relationships and reduce slow shared fluctuations, but significant short-lag interactions can still arise from monosynaptic coupling, polysynaptic pathways, common input, unobserved intermediate neurons, or state-dependent coordination among distributed populations (Smith and Kohn, 2008; Stark and Abeles, 2009; Jia et al., 2013, 2022); by the same logic, negative CCG peaks should not be equated directly with inhibitory synapses. Our conclusions therefore concern state-dependent reconfiguration of fast-timescale communication, not changes in anatomical wiring or direct causal links.

Second, locomotion is a coarse label for a richer behavioral and physiological state. Running covaries with arousal, neuromodulatory tone, eye movements, whisking, facial movements, and other uninstructed behaviors (Vinck et al., 2015; Reimer et al., 2014, 2016; Stringer et al., 2019; Musall et al., 2019; Akella et al., 2025). Our binary locomotion–stationary classification captures a robust and ethologically meaningful transition, but it does not identify which behavioral or neuromodulatory variables drive structured sparsification.

These constraints motivate several complementary experimental directions. First, causal tests would combine large-scale recordings with optogenetic perturbations to determine whether effective networks are reconfigured according to the same structured-sparsification framework. Second, future behavioral-state analyses could infer signal-transmission networks across a more continuous state manifold, incorporating running, pupil size, eye position, facial motion, body movement, and neuromodulatory signals, to clarify which dimensions of state drive the reconfiguration. Future studies could test whether structured sparsification is cell-type specific. Because locomotion modulates firing rates differently across cell types (Dipoppa et al., 2018; Polack et al., 2013), it may also reshape functional connectivity in a cell-type-dependent manner. Identifying the neuronal populations that drive these changes would clarify whether structured sparsification reflects a global network-state shift or selective reconfiguration of certain excitatory or inhibitory circuits. Moreover, the biological mechanisms underlying this structured sparsification remain unclear. Locomotion-induced changes in neural activity have been attributed to neuromodulatory systems (McGinley et al., 2015) and cell-type-specific modulation (Polack et al., 2013; Dipoppa et al., 2018). Whether these mechanisms can account for the observed structure of sparsification, and through what cellular or circuit-level processes, remains an open question. Addressing it will require careful cell type and neuromodulation monitoring, which is an important direction for future work.

In summary, structured sparsification is a candidate mechanism by which behavioral state reconfigures the architecture of cortical communication to improve coding. By linking experimentally inferred signal-transmission networks to signal-noise geometry and neural dynamics, our results provide a framework for understanding how network architecture shapes the capacity and timing of population codes.

## Methods

### Dataset

The data analyzed and discussed in this paper are part of the publicly released Allen Institute Brain Observatory Neuropixels dataset (Siegle et al., 2021). The Neuropixels project uses high-density extracellular probes to record neuronal spiking activity simultaneously from multiple regions of the mouse brain. In this work, we analyzed recordings obtained from six Neuropixels probes targeting six visual cortical areas (V1, LM, RL, AL, PM, AM) and two subcortical areas (LGN, LP). During data acquisition, mice were head-fixed but allowed to run freely on a wheel equipped with an encoder that continuously tracked running speed and distance. While recording, mice passively viewed six categories of visual stimuli: gray screens, full-field flashes, drifting gratings, static gratings, natural scenes, and natural movies. Trials were classified as locomotion if the mean running speed during stimulus presentation exceeded 1 cm/s, and as stationary otherwise. Sessions were excluded if trials in either state accounted for less than 20% of all trials. To balance behavioral conditions, we subsampled trials such that the number of trials in the two states was equal within each session. Only neurons with mean firing rates above 2 Hz during stimulus presentation in both states were included. Following these criteria, an average of 228 ± 61 units per mouse were retained for analysis.

### Statistics and data analyses

We used Python as the primary programming language for data analyses, together with analytical tools including SciPy, Pandas, scikit-learn and networkx. For pairwise comparisons between locomotion and stationary states measured across mice (*n* = 17 mice for drifting gratings, *n* = 24 mice for static gratings), the Wilcoxon signed-rank test was used for non-normally distributed metrics (e.g., network density, peak offset, peak width, modularity) and paired *t*-tests were used for metrics whose paired differences were approximately normally distributed (e.g., relative density changes between within-area and between-area connections, selectivity indices, distance ratio). Error bars, unless otherwise specified, represent standard error of the mean (SEM). For the feedforwardness analysis, error distributions were estimated by bootstrap resampling over mice (*n* = 20 repeats, 20% of mice per repeat), and significance was assessed using a paired *t*-test. Decoding performance was evaluated using multinomial logistic regression with repeated stratified 5-fold cross-validation (4 repeats). For modularity analysis, 50 surrogate networks per session were generated using a pair-preserving null model, and only modules containing at least four neurons were retained.

### Firing rate-matching control

To test whether locomotion-induced firing-rate modulation accounted for differences in network density, we performed a firing-rate-matching control. For each neuron, spikes were randomly subsampled from the behavioral state with the higher firing rate until its firing rate matched that in the lower-rate state. Thus, for most neurons, spikes were subsampled from locomotion trials to match the stationary firing rate, whereas for the small fraction of neurons with higher stationary firing rates, spikes were subsampled from stationary trials to match the locomotion firing rate. We then inferred signal-transmission networks from the rate-matched spike trains using jitter-corrected CCGs and compared network densities across behavioral states.

### Connectivity

We analyzed interactions between pairs of simultaneously recorded neurons by calculating the spike-train cross-correlogram (CCG) (Jia et al., 2022; Siegle et al., 2021; Smith and Kohn, 2008; Tang et al., 2024). For a pair of neurons with spike trains *X* and *Y*, the CCG at lag τ was defined as

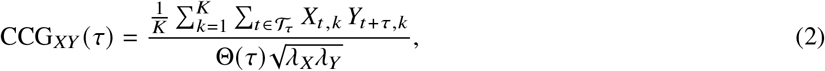

where *K* is the number of trials, *T* is the number of time bins in each trial, *X*_*t,k*_ and *Y*_*t,k*_ are the spike counts of the two neurons at time bin *t* on trial *k*, and *T*_τ_ = {*t*: 1 ≤ *t* ≤ *T*, 1 ≤ *t* +τ ≤ *T*} denotes the set of valid time bins for lag τ. The parameters λ_*X*_ and λ_*Y*_ are the mean firing rates of the two neurons. The factor Θ (τ) is the triangular correction for the number of overlapping time bins at lag τ.

The jitter-corrected CCG was computed by subtracting the expected CCG produced by spike trains with spike times randomly jittered within a jitter window (Harrison and Geman, 2009; Smith and Kohn, 2008). Jitter-corrected CCGs can be computed analytically (Harrison and Geman, 2009). For spike train *X*, let *W*_*s*_ index be the windows of length *L*, and let

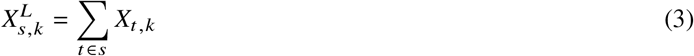

be the spike count in jitter window *s* on trial *k*. The across-trial mean count in that jitter window was

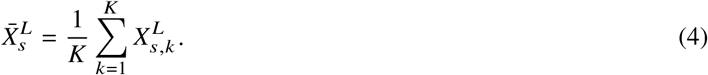

We normalized the binned spike counts as

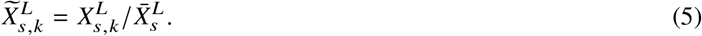

After expanding 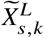 back to the original time resolution by assigning the same value to each time bin within window *s*, the PSTH-jitter mean for spike train *X* was

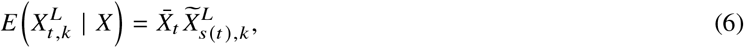

where *s*(*t*) denotes the jitter window containing time bin *t*, and 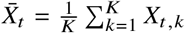. The same procedure was applied to spike train *Y*.

The jittered CCG was then computed as

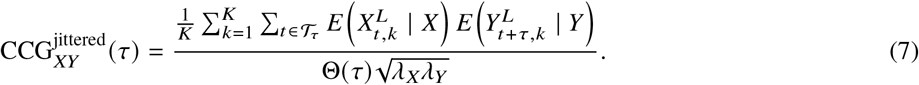

Finally, the jitter-corrected CCG was

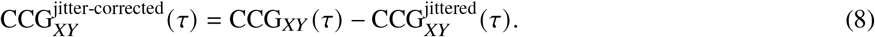

Consistent with our previous studies, we chose a jitter window of *L* = 25 ms. A positive sharp peak was deemed significant if the maximum of the jitter-corrected CCG amplitude within the 0–13 ms time window exceeded *μ* + 4σ. A negative sharp peak was deemed significant if the minimum of the jitter-corrected CCG amplitude within the same time window was lower than *μ* − 4σ. Here, *μ* and σ indicate the mean and standard deviation of jitter-corrected CCG values during [−100 ms, −50 ms] ∪ [50 ms, 100 ms].

### Directionality scores

The directionality score was computed following previous work (Siegle et al., 2021). For each pair of visual areas, we denoted the number of significant CCG connections from source area (area A) to target area (area B) as *C*_*BA*_ and the number of connections from target area (area B) to source area (area A) as *C*_*AB*_. The directionality score from area A to area B was defined as

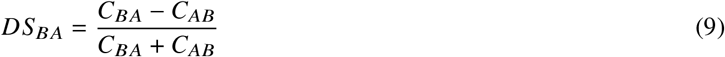

Thus, *DS*_*BA*_ > 0 indicates that there are more connections from area A to area B, while *DS*_*BA*_ < 0 indicates otherwise. We also computed a directionality score based on reachability. A target neuron was considered reachable from a source neuron if there was a directed path, consisting of one or more directed connections, from the source to the target. The number of reachable pairs from area A to area B was *R*_*BA*_, and the reachability-based directionality score was

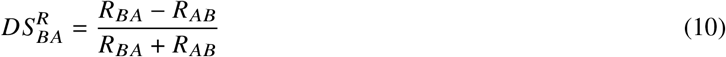

Canonical correlation analysis (CCA) was also used to quantify directional population interactions between brain areas (Semedo et al., 2022). For each ordered pair of visual areas, we computed the canonical correlation between population activity vector in area A *x*_*A*_ (*t*) and the time-lagged population activity vector in area B *x*_*B*_ (*t* +τ). We used the first canonical correlation as the interaction strength between the two areas. For each direction, we summed the first canonical correlations across positive lags to obtain the total lagged interaction strength from area *A* to area *B*,

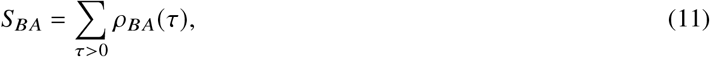

where *ρ*_*BA*_ (τ) is the first canonical correlation between activity in area *A* at time *t* and activity in area *B* at time *t* +τ. Similarly, *S*_*AB*_ denotes the summed lagged interaction strength in the opposite direction. The CCA-based directionality score was defined as

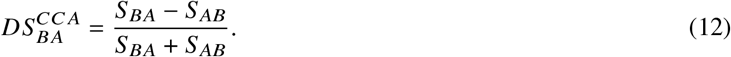

We computed this score separately for locomotion and stationary trials, using matched trial counts across behavioral states. Spike counts were extracted in 10-ms windows with 5-ms steps, and lags from 5 to 45 ms were evaluated for each ordered pair of visual areas.

### Network model

The network consisted of multiple layers representing different areas along the visual hierarchy. Neurons in each layer were arranged on a uniform grid covering a unit square Γ = [0, 1] × [0, 1]. Each layer contained *N* neurons, with 80% excitatory and 20% inhibitory. To simplify the representational landscape of image space, each neuron was tuned to a single angular feature representing the orientation of static gratings, randomly and uniformly sampled in [0, π]. The preferred orientation of neuron *i* was denoted by 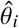. We assumed that the input from LGN can be modeled as

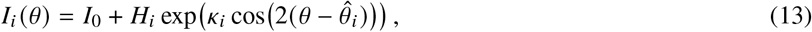

where *κ*_*i*_ controls the sharpness of this LGN input tuning. Unless otherwise stated, we set *I*_0_ = 1.0, *H*_*i*_ = 1, and *κ*_*i*_ = 1.5. Neuronal activity followed a rate equation:

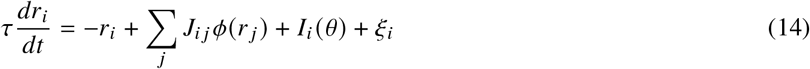

where τ = 10 is the membrane time constant, *J*_*ij*_ is the connection from neuron *j* to neuron *i*, and *ξ*_*i*_ is an independent Gaussian noise with 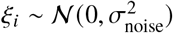 and σ_noise_ = 1.5. The transfer function was *ϕ*(*x*) = max (0, *x*). Synaptic strengths were set to *J*_*ee*_ = 0.06, *J*_*ie*_ = 0.06, *J*_*ei*_ = −0.24, and *J*_*ii*_ = −0.28.

### Distance-dependent connectivity

The total connection density within layer *n* is *d*_*nn*_. The cross-layer connection density from source layer *m* to target layer *n* is denoted *d*_*nm*_. Within each layer, the connection probability from neuron *j* to neuron *i* is distance-dependent:

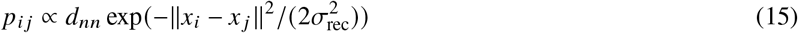

where *x*_*i*_ and *x*_*j*_ denote the spatial coordinates of neurons *i* and *j*, respectively, and σ_rec_ controls the spatial scale of recurrent connectivity. Feedforward connections from layer 1 to layer 2 also followed the same distance-dependent rule, but with a different spatial scale parameter σ_ffwd_. Following the convention above, for a connection from neuron *j* in source layer *m* to neuron *i* in target layer *n*, the connection probability is

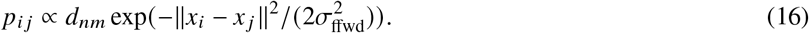

### Feature-specific connectivity

Feature-specific connectivity was achieved by making the connection probability depend on feature similarity:

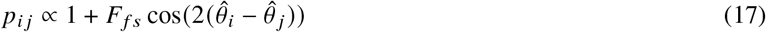

with *F*_*fs*_ controlling the strength of feature specificity (0 ≤*F*_*fs*_ ≤1). 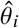 and 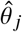 are the preferred orientation of neuron i and neuron j, respectively.

Taken together, the connection probability from neuron *j* in source layer *m* to neuron *i* in target layer *n* is

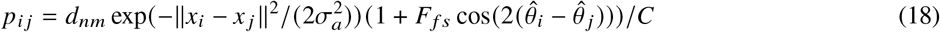

where *C* is the normalization term.

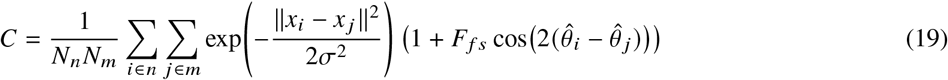

*N*_*n*_ and *N*_*m*_ denote the numbers of neurons in layers *n* and *m*, respectively, both of which equal *N*. σ_*a*_ = σ_rec_ when *n* = *m*, and σ_*a*_ = σ_ffwd_ otherwise.

### Increased number of input projections to weakly selective neurons

To model the observation that highly selective neurons receive fewer recurrent inputs whereas non-selective neurons receive more recurrent inputs, we introduced a hyperparameter *β*. Specifically, each neuron’s *κ*_*i*_ (see equation (13)) was independently sampled from a gamma distribution,

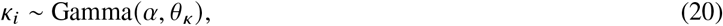

where we used shape *α* = 0.7 and scale *θ*_*k*_ = 1.0.

Let 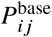 denote the baseline probability of a recurrent connection from presynaptic neuron *j* to postsynaptic neuron *i*, after incorporating the spatial and feature-specific connectivity terms. We defined a postsynaptic targeting factor based on the *κ*_*i*_ value:

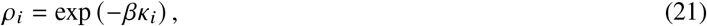

*β* = 0 indicate that all neurons have the same number of input projections, while higher *β* corresponds to increased the number of inputs to weakly selective neurons and reduced number of inputs to highly selective neurons. The unnormalized biased connection probability from neuron *j* to neuron *i* is

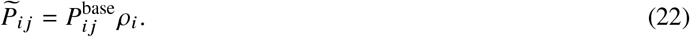

For each presynaptic neuron *j*, we then normalized the probabilities across all valid postsynaptic targets to preserve the original expected out-degree of neuron *j* :

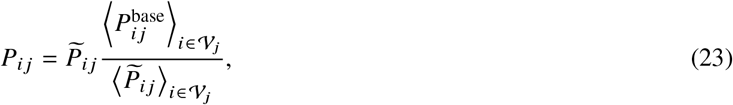

where *V*_*j*_ denotes the valid postsynaptic targets for presynaptic neuron *j*, excluding self-connections for recurrent connectivity and ⟨ ⟩_∈V*j i*_ denotes the average over the valid postsynaptic targets of neuron j.

Thus, *β* = 0 leaves the baseline connection probabilities unchanged. For *β* > 0, neurons with larger *κ*_*i*_ have smaller targeting factors *ρ*_*i*_ and therefore receive fewer recurrent inputs on average, whereas neurons with smaller *κ*_*i*_ receive more recurrent inputs. The targeting factor modulates only the probability that a postsynaptic neuron is targeted; it does not directly modify synaptic weights. After normalization, probabilities were clipped to the valid range [0, 1] when necessary.

### Feedforwardness of inter-area connectivity

We introduced a feedforwardness parameter *f* that controls the the fraction of feedforward connections.

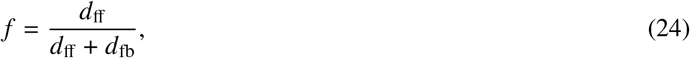

where *d*_ff_ and *d*_fb_ denote the densities of feedforward and feedback inter-area connections, respectively. Thus, *f* = 1 corresponds to a purely feedforward architecture, *f* = 0.5 to balanced feedforward and feedback connectivity, and *f* = 0 to a purely feedback architecture. Feedforward and feedback projections used the same spatial and feature-dependent connectivity rules; *f* controls only projection direction.

### Simulations

All simulations were performed by numerically integrating the rate equations with a forward Euler method. For each parameter setting, results were averaged across 20 independently generated network realizations. We used nine stimulus orientations ranging uniformly from 0 to π. Forty trials were simulated per stimulus orientation with independent Gaussian noise on each trial and time step. Activity was initialized at zero and simulated for 200 time steps; the average response over this period was used as the trial response. For time-resolved analyses of trajectory geometry and decoding latency, simulations additionally included a 50-time-step prestimulus baseline and a 250-time-step poststimulus period.

### Intrinsic dimensionality

Intrinsic dimensionality was computed as the participation ratio of eigenvalues of the noise covariance matrix. Mathematically, let λ_*i*_ be the *i*th eigenvalue of the noise covariance matrix Σ. Intrinsic dimensionality is

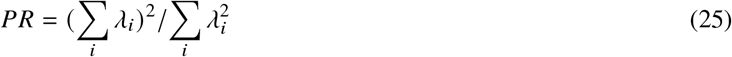

### Linear Fisher information

To quantify the stimulus information available to an optimal *linear* decoder, we computed the linear Fisher information around a reference stimulus value *θ*_0_ as

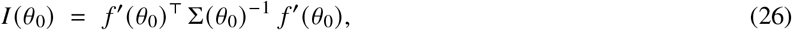

where *f* (*θ*) = E[*r* | *θ*] ∈ ℝ^*N*^ denotes the mean population response (tuning curve) and 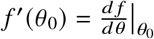 is the vector of tuning-curve derivatives with respect to the stimulus parameter. Σ (*θ*_0_) ∈ ℝ^*N* ×*N*^ is the noise covariance of the trial-to-trial response variability around *θ*_0_.

In practice, we computed the generalized linear Fisher information (Kafashan et al., 2021). We estimated *f*′(*θ*_0_) and Σ(*θ*_0_) by symmetric finite differences using two nearby stimulus values *θ*_1,2_ = *θ*_0_ ± *δθ*/2:

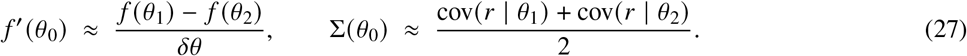

This approximation treats the covariance as locally constant over the interval *δθ* and reduces bias relative to a one-sided difference. We then bias-corrected the estimated linear Fisher information (Kanitscheider et al., 2015a):

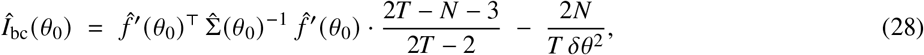

where *T* is the number of trials and *N* is the number of neurons.

### Eigenmode and geometric interpretation of linear Fisher information

To provide a geometrically interpretable decomposition, we expressed linear Fisher information in the eigenbasis of the covariance. For the noise covariance matrix Σ(*θ*_0_), let

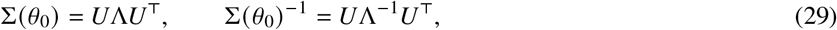

where *U* = [*u*_1_, …, *u*_*N*_] contains the eigenvectors and Λ = diag (λ_1_, …, λ_*N*_) contains the eigenvalues (noise variances along the principal axes). Expressing the signal direction in this eigenbasis gives

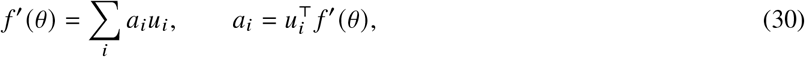

so that Fisher information can be written as

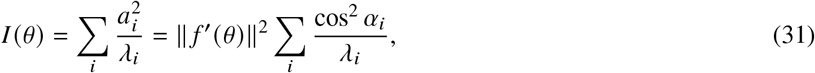

where *α*_*i*_ is the angle between *f*′ (*θ*) and the *i*th covariance eigenvector. Thus, linear Fisher information is determined by three geometric factors: the magnitude of the signal, ∥*f*′ (*θ*) ∥; the magnitude of noise along each covariance axis, λ_*i*_; and the alignment of the signal with high- or low-variance noise directions, quantified by cos^2^ *α*_*i*_.

### Modularity

The modularity for directed networks (Leicht and Newman, 2008) is defined as

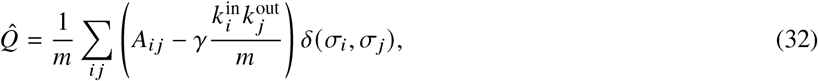

where *A* is the adjacency matrix of the network, *m* is the number of edges, and 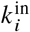 and 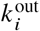 represent the in-degree and out-degree of node *i*, respectively. *δ* is the Kronecker delta function, and σ_*i*_ denotes the community assignment of node *i. δ* (σ_*i*_, σ_*j*_) = 1 if and only if nodes *i* and *j* are assigned to the same community. *γ* is a resolution parameter that controls the size of modules. We ignored connection strength as well as the sign of connections by only considering the existence of edges. The Louvain method is a community-detection algorithm for partitioning networks into groups of nodes with dense connections within groups and sparse connections between groups (Blondel et al., 2008). The algorithm uses the original modularity 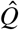 as a quality function to optimize the partitioning of the network. The Louvain method operates through a series of iterative steps that merge neighboring modules to maximize modularity gain until a locally optimal partition is reached. The algorithm follows a bottom-up approach, starting from single-node modules and iteratively merging modules to form larger ones. The Louvain method was used to compute modularity.

To determine the resolution parameter for module analysis, we obtained a modularity-difference curve by varying *γ* and computing the difference between the modularity of empirical and surrogate networks generated by the pair-preserving model, and then selected the value of *γ* that maximized this difference (Harris et al., 2019; Tang et al., 2024). The pair-preserving model randomizes the network while preserving network size, density, degree distribution, and neuron pair distribution (Milo et al., 2002; Tang et al., 2024). This procedure yielded the module partition that deviated most from the reference model. We used the *Z*-score of modularity to quantify the strength of modular organization relative to a reference model. The *Z*-score of modularity is defined as

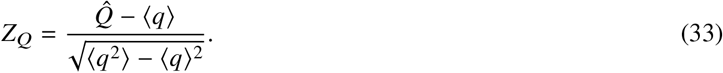

Here, *q* denotes the modularity obtained from one realization of the reference model. We randomly generated 50 surrogate networks for each session. Only modules with a size of at least four neurons were included in subsequent analyses to eliminate noise contributions from isolated single neurons, pairs, and triplets.

### Selectivity index

In total, eight different directions of drifting gratings, evenly spaced in [0, 2π), and six different orientations of static gratings, evenly spaced in [0, π), were presented to the mice. We calculated each neuron’s direction selectivity index (DSI) from its drifting-grating tuning curve and its orientation selectivity index (OSI) from its static-grating tuning curve as follows:

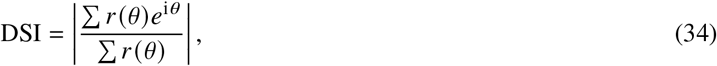

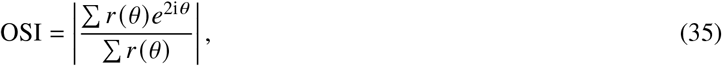

where *θ* is the stimulus direction or orientation and *r* (*θ*) is the mean neuronal response for a given stimulus parameter *θ*.

### Selectivity-enhancing input projections

Consider a population of neurons responding to external stimuli parameterized by their orientation, *θ*. The response of neuron *m* to a stimulus with orientation *θ* is denoted by *r*_*m*_ (*θ*). When neuron *m* receives an input projection from neuron *n*, its modified response is modeled as

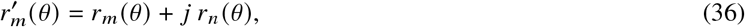

where *j* denotes the effective coupling strength. We define the response-weighted coupling strength as

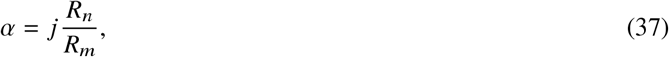

where *R*_*m*_ and *R*_*n*_ denote the response amplitudes of neurons *m* and *n*, respectively. We also define the difference between their preferred orientations as

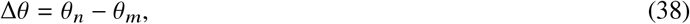

where *θ*_*m*_ and *θ*_*n*_ are the preferred orientations of neurons *m* and *n*, respectively. The modified orientation selectivity of neuron *m* is then given by

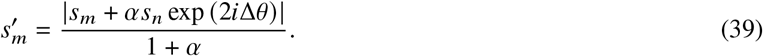

Equivalently, the exact post-input selectivity is

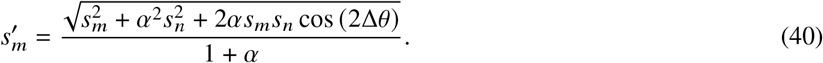

For weak coupling,

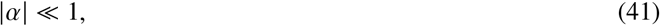

a first-order expansion of Eq. (40) in *α* gives

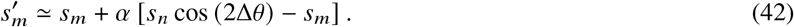

Therefore, the first-order change in the selectivity of neuron *m* is

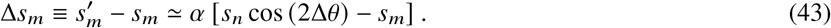

The excitatory input projection enhances the selectivity of neuron m when

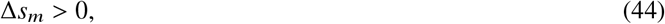

or, equivalently,

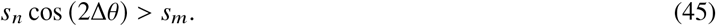

This condition can be satisfied only if the presynaptic neuron is more selective than the postsynaptic neuron:

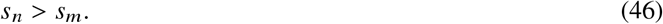

When *s*_*n*_ > *s*_*m*_, their preferred orientations must also be sufficiently aligned:

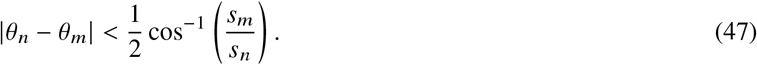

We refer to input projections that satisfy these selectivity and alignment conditions as selectivity-enhancing projections. An inhibitory input projection enhances selectivity when

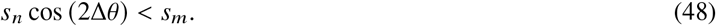

### Distance ratio

The distance ratio was computed as previously reported (Horrocks et al., 2024). We denote the neural response at time *t* as *r*_*t*_, which is an *N* × 1 vector, with *N* being the number of neurons. *R* = [*r*_1_, …, *r*_*T*_]. We applied principal component analysis to *R* and retained the principal dimensions that explain 90% of total variance, yielding a lower-dimensional representation matrix *X* of size *m* × *T*, where *m* is the number of dimensions. *x*_*t*_ is the position of the neural state in *m*-dimensional neural space at time *t*. To quantify how direct the trial-averaged population trajectory was as it moved between stimulus onset and stimulus offset, we computed a distance ratio. This ratio is the cumulative path length traveled by the trajectory between the two endpoints, divided by the Euclidean distance between those same endpoints.

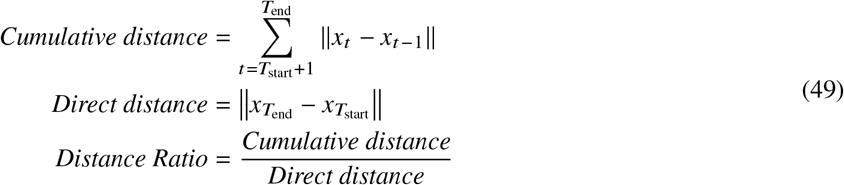

where *T*_start_ is the time of stimulus onset and *T*_end_ is the time of stimulus offset.

### Tangling index

We calculated population tangling using previously described methods (Horrocks et al., 2024; Russo et al., 2018). Trajectory tangling was quantified as

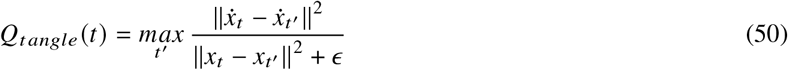

where *x*_*t*_ is the PCA-decomposed neural state in *m*-dimensional neural space at time *t* (as described in the Distance ratio section), which is an *m* × 1 vector, with *m* being the number of dimensions. 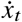 is the temporal derivative of *x*_*t*_. We computed the derivative of *x*_*t*_ as 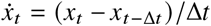, with Δ*t* = 1 ms. ∥ · ∥ is the Euclidean norm, and ϵ is a small constant that prevents division by zero. Neural tangling *Q*_*tangle*_ *t* is large when the neural population state at time *t* closely matches a state observed at another time *t*′, yet the associated state velocities (time derivatives) differ substantially. To obtain a temporally local estimate, we constrained the search for *t*′ to within ± 100 ms of *t*. This avoids spuriously large tangling driven by temporally distant epochs.

### Peak width

To characterize the temporal precision of pairwise coupling between neuronal signals, we quantified the peak width of the cross-correlogram (CCG) using a smoothed half-amplitude approach. For each significant sharp CCG peak, we first baseline-corrected the trace by subtracting the mean CCG value computed over the baseline period (time lags from −100 to −50 ms and 50 to 100 ms). The baseline-corrected traces were then smoothed using a Savitzky–Golay filter (Gorry, 1991) (window length = 11 ms, polynomial order = 3) to suppress high-frequency noise while preserving the overall peak shape. Positive and negative peaks were treated separately. For positive connections, we identified the time point of the maximum correlation and used one-half of the positive peak amplitude as the width threshold. For negative peaks, we identified the time point of the minimum correlation and used one-half of the absolute negative peak amplitude as the width threshold. The left and right boundaries were defined as the nearest time points on either side of the peak extremum where the smoothed trace crossed the corresponding half-amplitude threshold. The peak width was then quantified as the temporal distance (in time bins) between these two crossing points.

### Population decoding analysis

To quantify stimulus information across behavioral states, we trained multinomial logistic regression classifiers on neuronal firing rates during stimulus presentation. Class labels corresponded to unique stimulus orientation and spatial frequency combinations for static gratings. And labels corresponded to direction and temporal frequency combinations for drifting gratings. Firing rates were z-scored across trials, and decoding performance was evaluated separately for each behavioral state using repeated stratified 5-fold cross-validation with four repeats. For decoding-latency analyses, firing rates were computed in 20-ms sliding windows with 2-ms steps. Decoding latency was defined as the first time window in which accuracy exceeded the 95th percentile of the chance distribution.

## Data availability

Data is freely accessible via the Allen SDK (Siegle et al., 2021) (Details at https://portal.brainmap.org/circuits-behavior/visual-coding-neuropixels). Source data for all main figures are provided with this paper.

## Code availability

Code for analyses in the manuscript and generation of figures are available from the repository: https://github.com/JiajZhu/Structured_sparsification

## Acknowledgements

This work was supported by funding from the Tsinghua-Peking Center for Life Sciences, National Natural Science Foundation of China (92370116), the Beijing Major Science and Technology Project (No. Z251100008125058 to X.J.), Tsinghua University Initiative Scientific Research Program, and Tsinghua University Dushi Program. We thank Disheng Tang, Zimoren Zhang, Ying Zhou and Fan Ye for helpful discussions, and Wenhao Zhang and Gabriel Koch Ocker for helpful feedback on the manuscript.

## Author contributions

J.Z. and X.J. conceived the study and designed the analyses. J.Z. performed the data analysis, developed the computational models, generated the figures, and drafted the manuscript. X.J. supervised the project, acquired funding, and provided conceptual and methodological guidance. J.Z. and X.J. interpreted the results, revised the manuscript, and approved the final manuscript.

## Extended Data Figures

**Extended Data Fig. 1.**
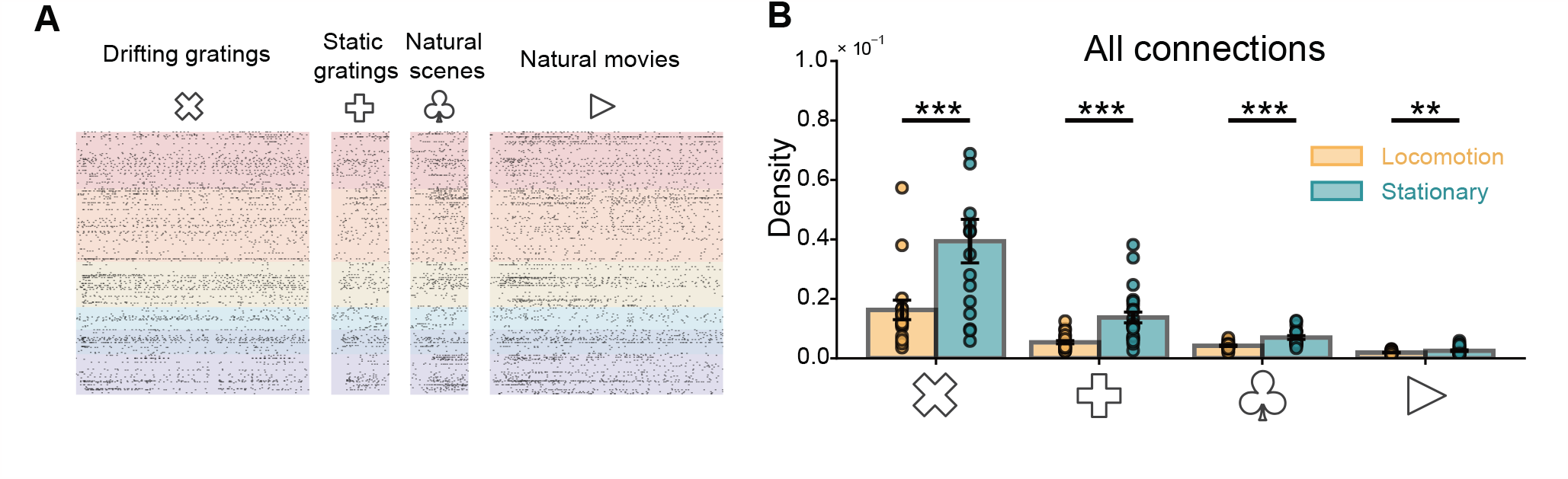
Locomotion-induced sparsification is robust to stimulus type. **A**, Example raster plots from mouse visual cortex during presentation of drifting gratings, static gratings, natural scenes, and natural movies. **B**, Network density across four visual stimulus types at a confidence threshold of 4. Points represent individual mice (Wilcoxon signed-rank test, from left to right *p* = 3.05 × 10^−5^, 4.77 × 10^−7^, 2.67 × 10^−6^, 2.64 × 10^−3^; *n* = 17, 24, 19, 24 mice).

**Extended Data Fig. 2.**
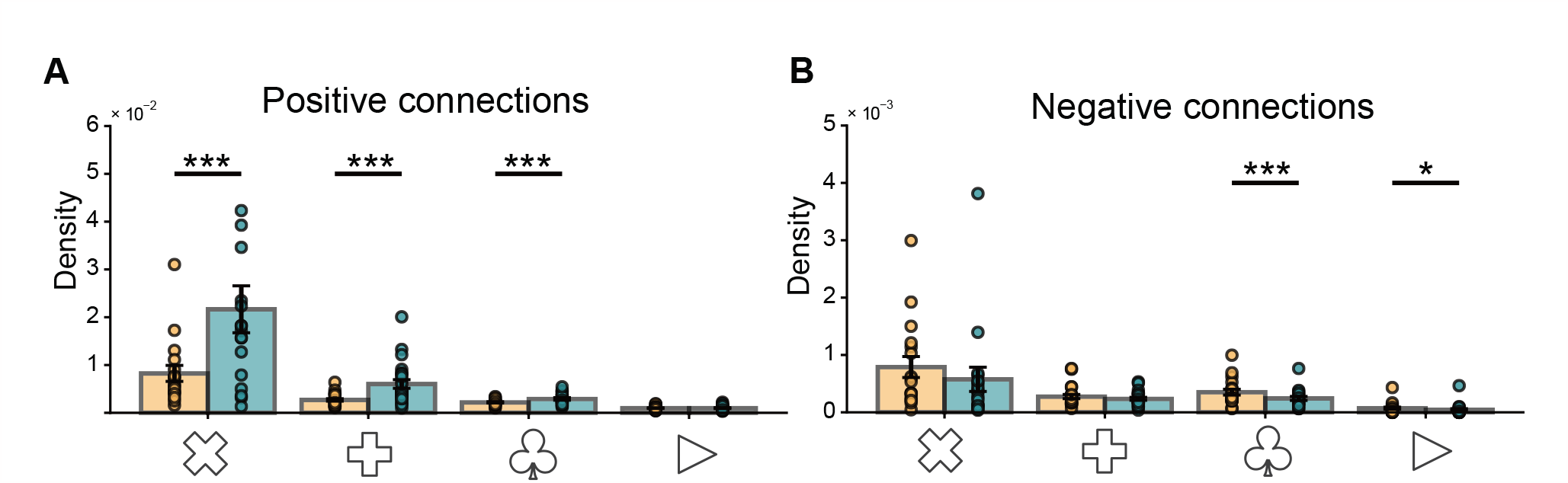
Sparsification in positive and negative connections. **A**, Density of positive connections across four visual stimulus types. Points represent individual mice (Wilcoxon signed-rank test, from left to right *p* = 3.05 ×10^−5^, 4.76 ×10^−7^, 3.81 ×10^−6^, 1.35×10^−1^; *n* = 17, 24, 19, 24 mice). **B**, Density of negative connections across four visual stimulus types (Wilcoxon signed-rank test, from left to right *p* = 6.00 ×10^−2^, 3.34 ×10^−1^, 3.77 ×10^−6^, 1.47 ×10^−2^; *n* = 17, 24, 19, 24 mice). Density of negative connections was not significantly different across states for drifting gratings and static gratings, and was higher during locomotion for natural scenes and natural movies.

**Extended Data Fig. 3.**
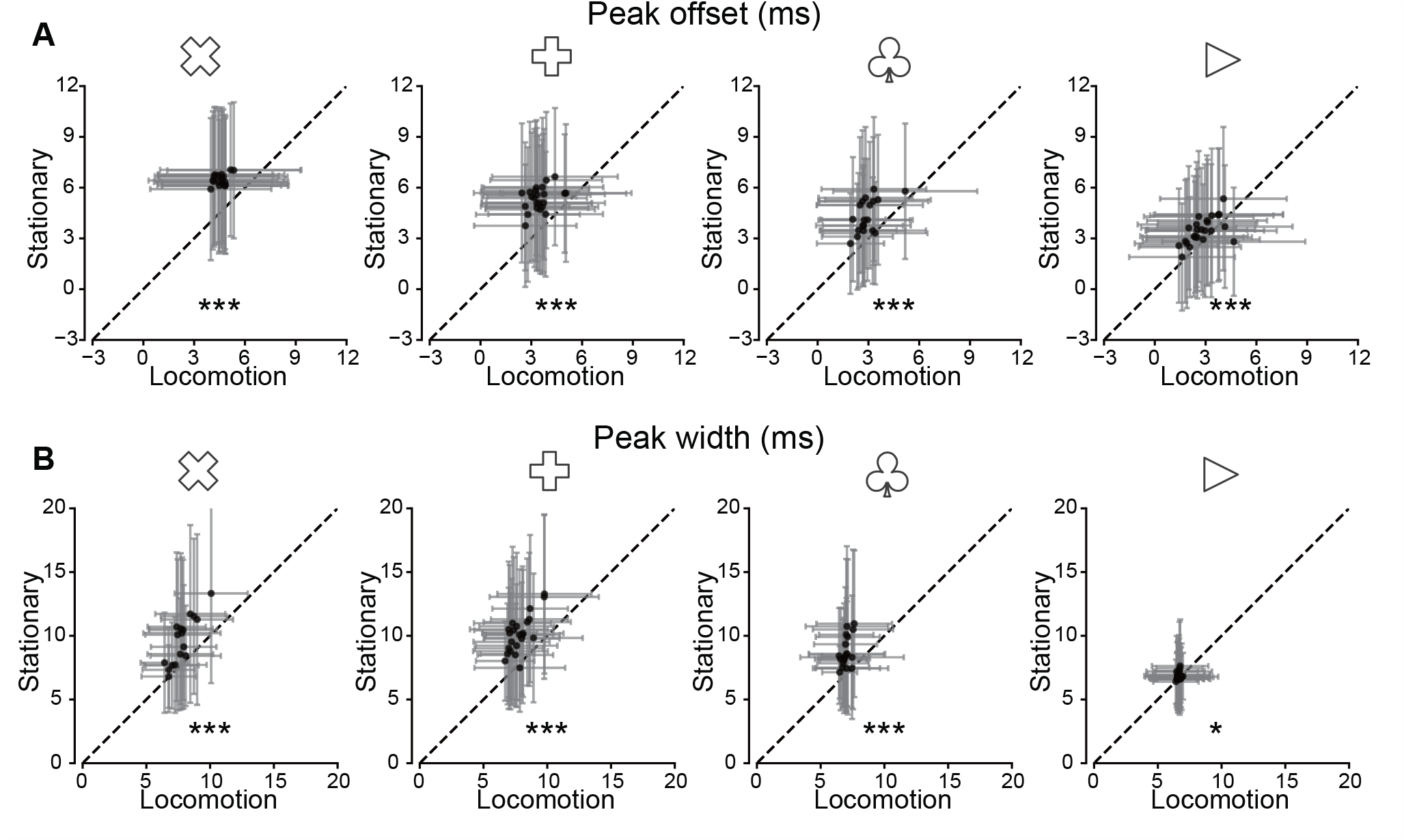
Offset and width of significant CCG peaks. **A**, Peak offset across four visual stimulus types. Points represent individual mice. Error bars indicate STD. (Wilcoxon signed-rank test, from left to right *p* = 1.5 ×10^−5^, 2.4 ×10^−7^, 7.6 ×10^−6^, 2.1 ×10^−4^; *n* = 17, 24, 19, 24 mice). **B**, Peak width across four visual stimulus types. Points represent individual mice. Error bars indicate STD. (Wilcoxon signed-rank test, from left to right *p* = 6.0 × 10^−4^, 4.8 × 10^−7^, 7.61 × 10^−6^, 1.2 × 10^−2^; *n* = 17, 24, 19, 24 mice).

**Extended Data Fig. 4.**
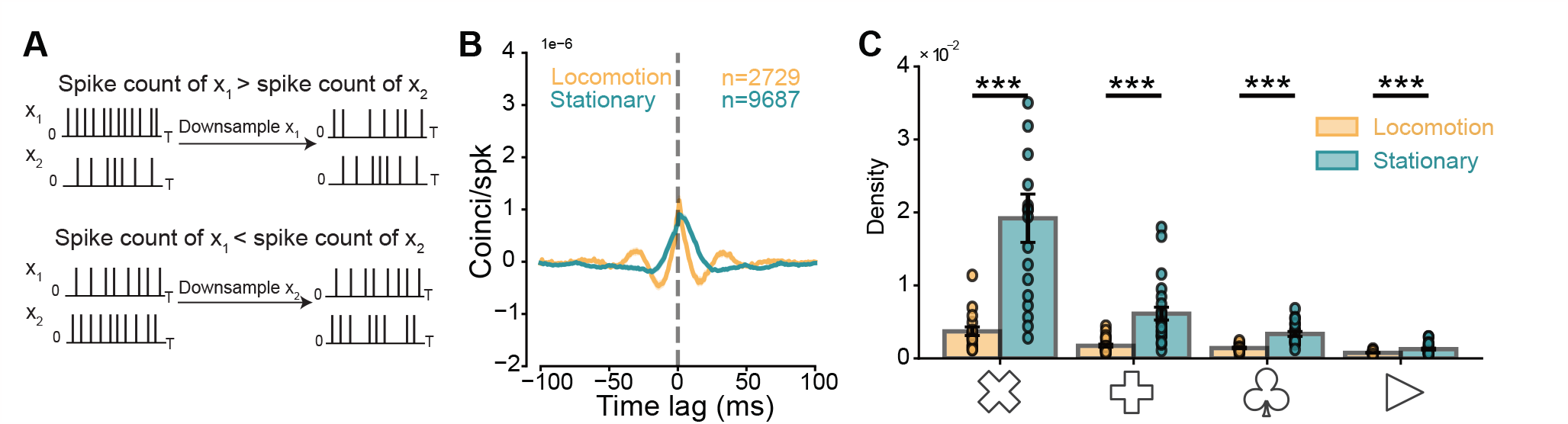
Controlling for firing rate does not eliminate the density reduction. **A**, Schematic illustration of spike downsampling. **B**, CCG trace of downsampled spikes with the same pairs in Fig. 1I. **C**, Network density inferred from downsampled spikes across four visual stimulus types. Points represent individual mice. Error bars indicate SEM. (Wilcoxon signed-rank test, from left to right *p* = 1.52 ×10^−5^, 2.38 ×10^−7^, 3.81 ×10^−6^, 3.65 ×10^−5^; *n* = 17, 24, 19, 24 mice). Across stimulus types, firing rates before rate matching during locomotion and stationary: drifting gratings, 8.511 ± 0.919 vs. 5.795 ± 0.623; static gratings, 9.098 ± 0.788 vs. 6.951 ± 0.594; natural scenes, 9.743 ± 0.711 vs. 7.358 ± 0.655; and natural movies, 8.784 ± 0.911 vs. 6.721 ± 0.885. After rate matching, firing rates were 5.470 ± 0.593, 6.511 ± 0.610, 6.938 ± 0.619, and 6.131 ± 0.749 for drifting gratings, static gratings, natural scenes, and natural movies, respectively.

**Extended Data Fig. 5.**
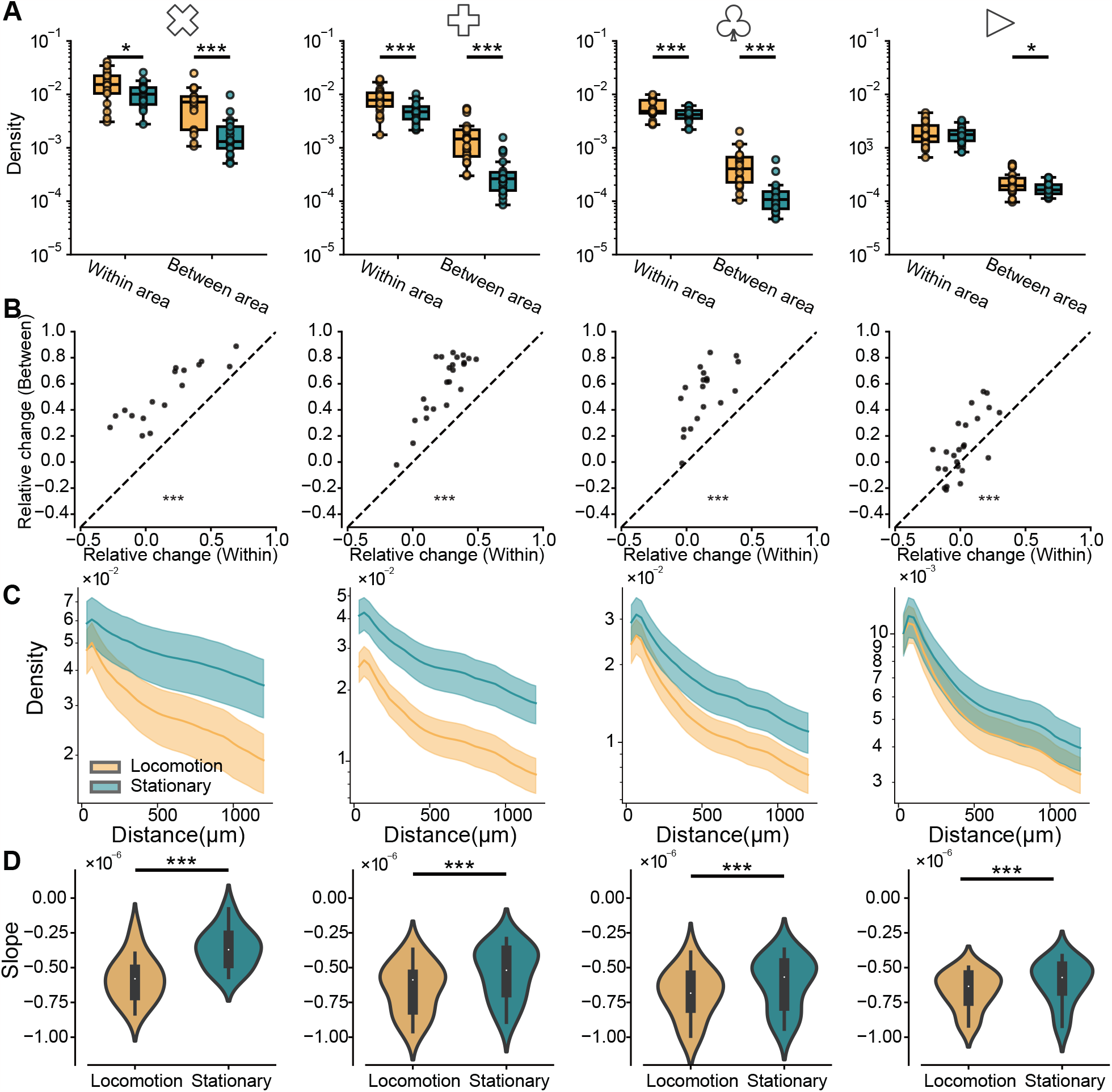
The anatomical pattern of sparsification is consistent across stimulus types. **A**, Within-area and between-area network density across four visual stimulus types. Box plots show the median (center line), interquartile range (box), and whiskers extending to 1.5 ×IQR. Points represent individual mice. (Wilcoxon signed-rank test, from left to right, within area: *p* = 1.74 ×10^−2^, 1.19 ×10^−6^, 6.45 ×10^−4^, 0.56; between area: *p* = 1.52 ×10^−5^, 4.76 ×10^−7^, 7.62 ×10^−6^, 0.03; *n* = 17, 24, 19, 24 mice). **B**, Relative change in network density across four visual stimulus types. Points represent individual mice. (Wilcoxon signed-rank test, from left to right, *p* = 1.52 ×10^−5^, 2.38 ×10^−7^, 3.81 ×10^−6^, 8.71 ×10^−4^; *n* = 17, 24, 19, 24 mice). **C**, Spatial decay of network density across four visual stimulus types. **D**, Slopes of the spatial decay of network density across four visual stimulus types. (Paired t-test, from left to right, *p* = 5.19 ×10^−7^, 2.31 ×10^−7^, 4.88 ×10^−4^, 7.70 ×10^−4^; *n* = 17, 24, 19, 24 mice).

**Extended Data Fig. 6.**
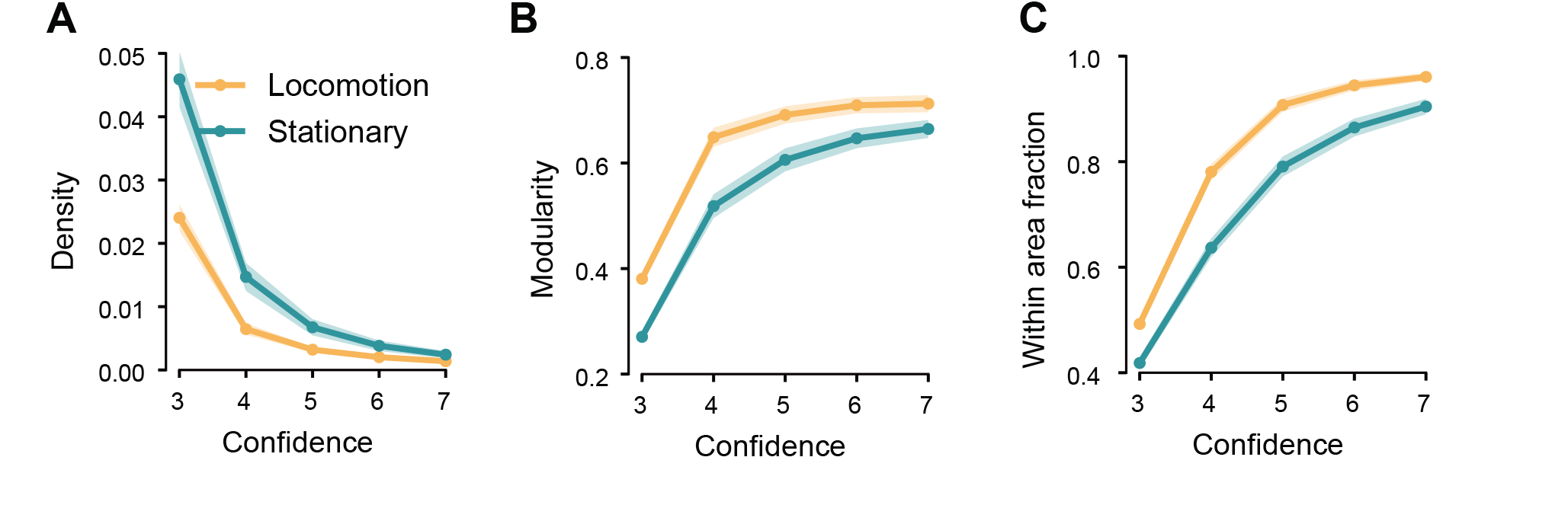
Structured sparsification across significant-peak thresholds. **A–C**, Density (**A**), modularity (**B**), and within-area fraction (**C**) across significant-peak thresholds. Shaded areas indicate SEM (*n* = 84 mice).

**Extended Data Fig. 7.**
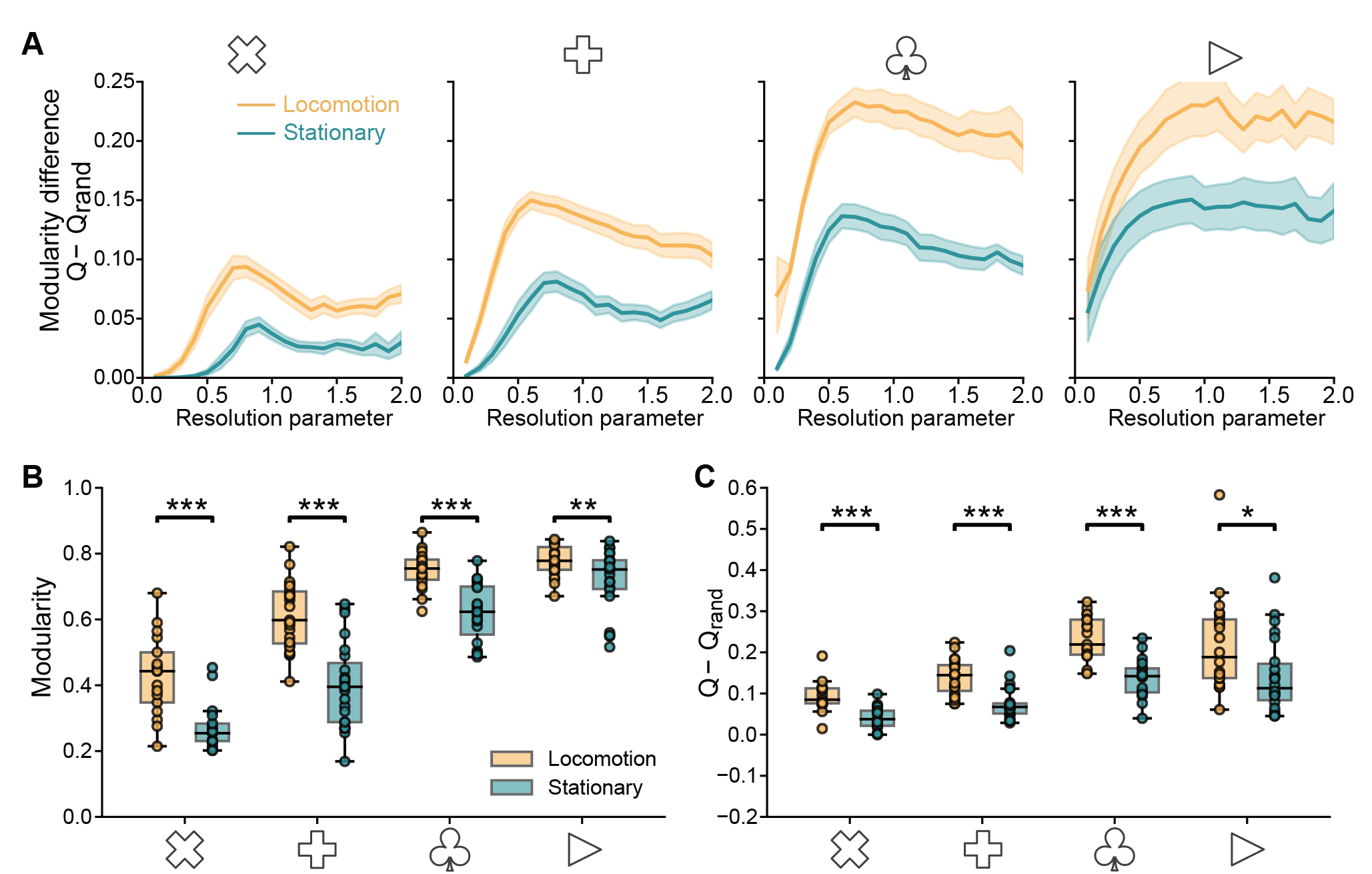
Network modularity across stimulus types. **A**, Relationship between the modularity difference and the resolution parameter. Shaded areas indicate SEM. (*n* = 17, 24, 19, 24 mice). **B**, Network modularity across four visual stimulus types. Box plots show the median (center line), interquartile range (box), and whiskers extending to 1.5× IQR. (Paired t-test, from left to right, *p* = 1.50 × 10^−5^, 4.15 × 10^−10^, 2.86 × 10^−6^, 7.45 × 10^−3^; *n* = 17, 24, 19, 24 mice). **C**, Network modularity difference (*Q* − *Q*_*rand*_) across four visual stimulus types. Box plots show the median (center line), interquartile range (box), and whiskers extending to 1.5× IQR. (Paired t-test, from left to right, *p* = 1.42 × 10^−6^, 1.05 × 10^−9^, 8.64 × 10^−6^, 1.22 × 10^−2^; *n* = 17, 24, 19, 24 mice).

**Extended Data Fig. 8.**
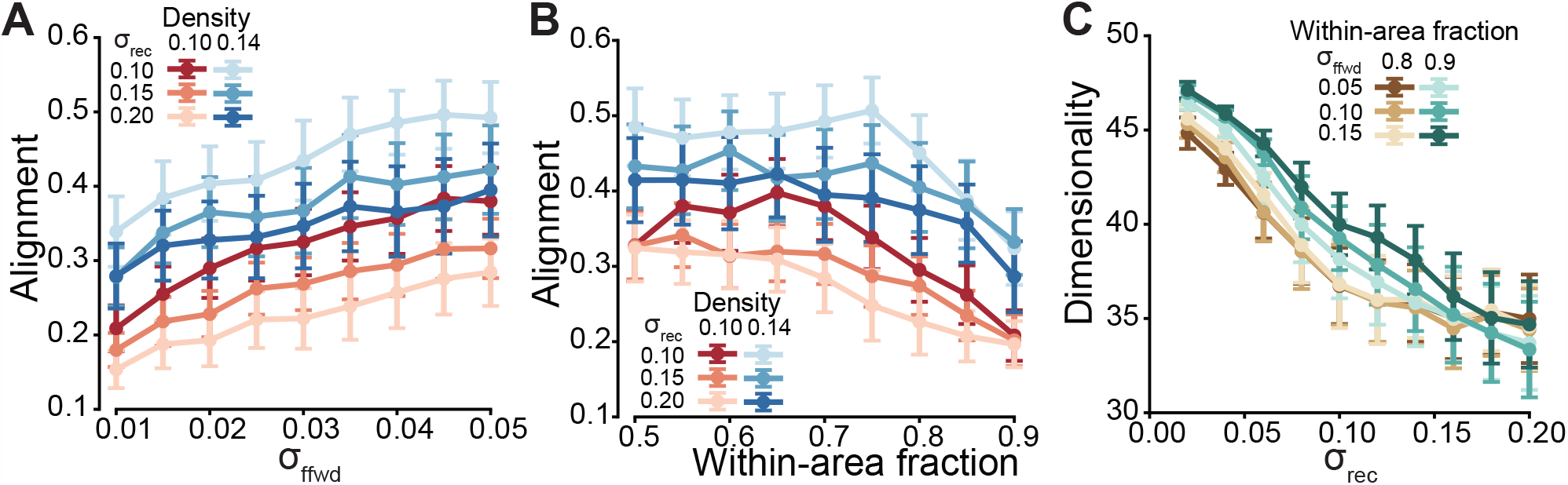
Spatial scale of connectivity shapes noise geometry. Network simulations showed that the spatial organization of recurrent and feedforward connectivity systematically reshaped the geometry of population variability across different choices of parameters. **A**, Increasing the spatial scale of feedforward connectivity, σ_ffwd_, increased signal–noise alignment, indicating stronger overlap between stimulus-driven signal directions and dominant noise-covariance modes. This effect was observed across different connection densities and recurrent spatial scales, σ_rec_. **B**, Increasing the fraction of within-area connections reduced signal–noise alignment across different densities and recurrent spatial scales. **C**, Increasing the recurrent spatial scale, σ_rec_, reduced intrinsic dimensionality, consistent with broader recurrent coupling concentrating variability into fewer dominant modes. Points and error bars indicate mean ± SEM across 20 simulation repeats.

**Extended Data Fig. 9.**
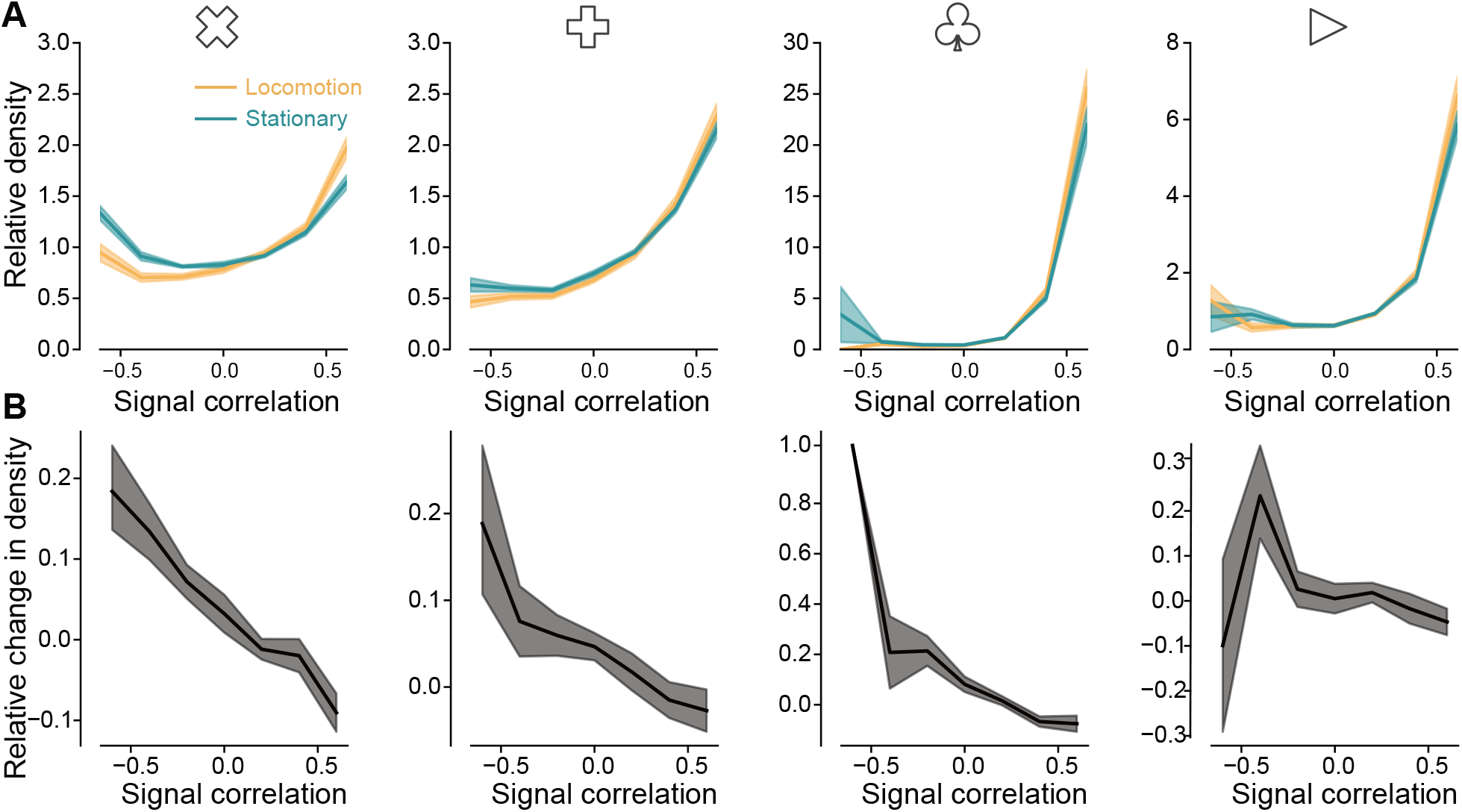
Feature-specific connectivity using signal correlation as a measure of feature similarity. **A**, Relationship between signal correlation and relative density across four visual stimulus types. **B**, Relationship between signal correlation and the relative change in density across four visual stimulus types. Shaded areas indicate SEM. (From left to right, *n* = 17, 24, 19, 24 mice).

**Extended Data Fig. 10.**
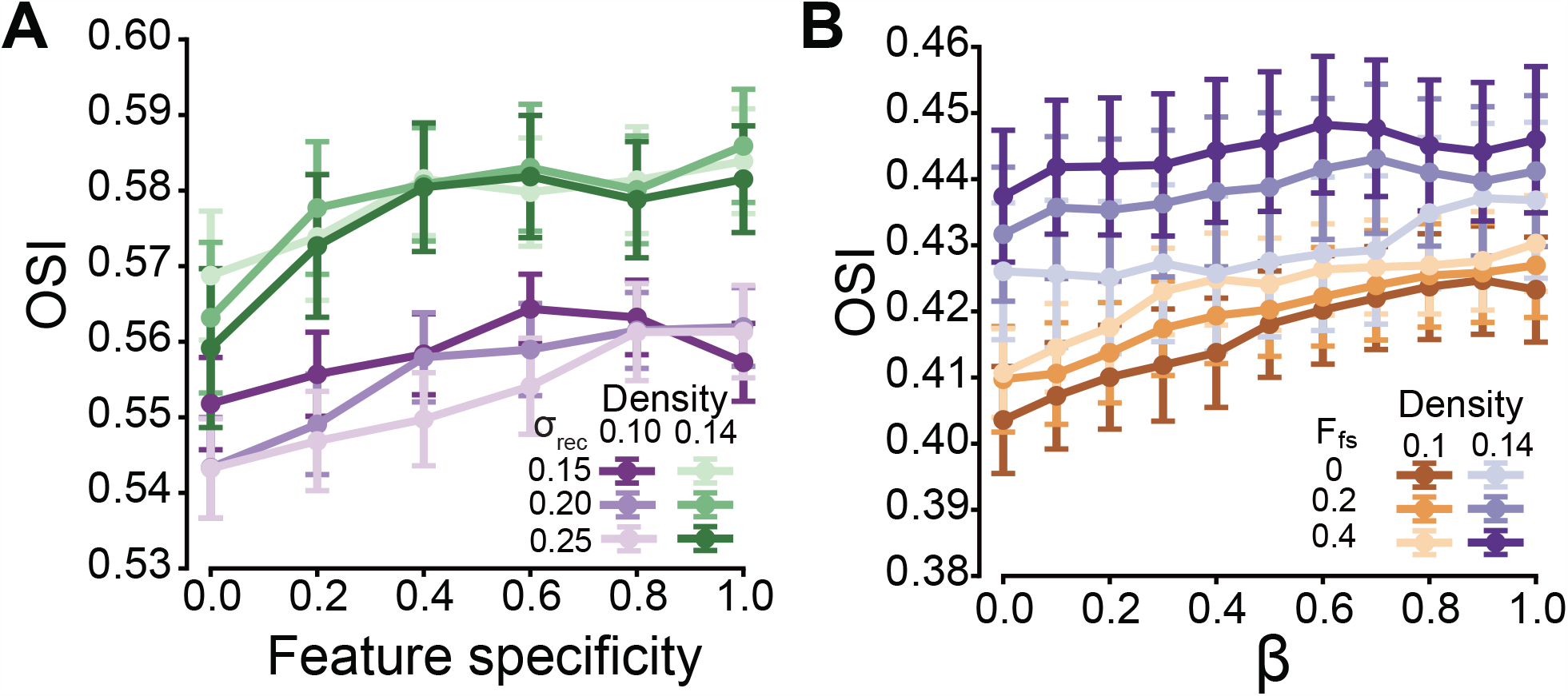
Structured sparsification increases selectivity. **A**, Selectivity increases as feature specificity increases, under different densities and spatial scales of recurrent connections. **B**, Selectivity increases as *β* increases, under different densities and feature specificity values. Points and error bars indicate mean ± SEM across 20 simulation repeats.

**Extended Data Fig. 11.**
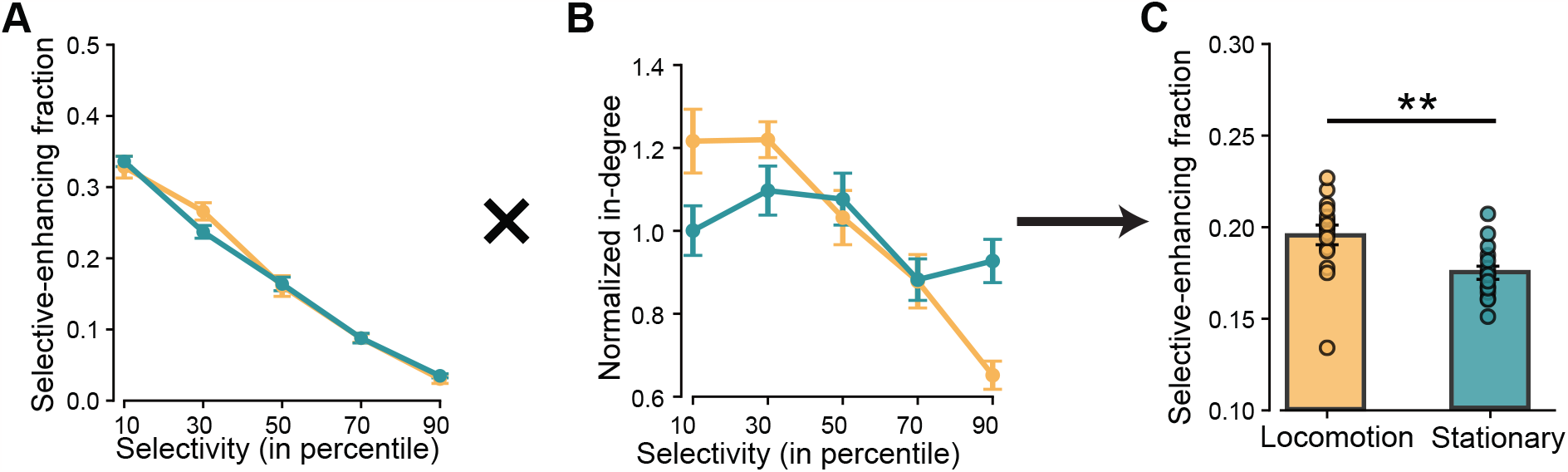
Fraction of selective-enhancing input projections. **A**, Fraction of selective-enhancing input projections across neuron populations with different selectivity. **B**, Normalized in-degree of neurons with different selectivity levels (percentile). Error bars indicate SEM (*n* = 17 mice). **C**, Overall fraction of selective-enhancing input projections. The overall selective-enhancing fraction was calculated by weighting each subpopulation’s selective-enhancing fraction by its in-degree. Error bars indicate SEM (Wilcoxon signed-rank test, *p* = 0.0021, *n* = 17 mice).

**Extended Data Fig. 12.**
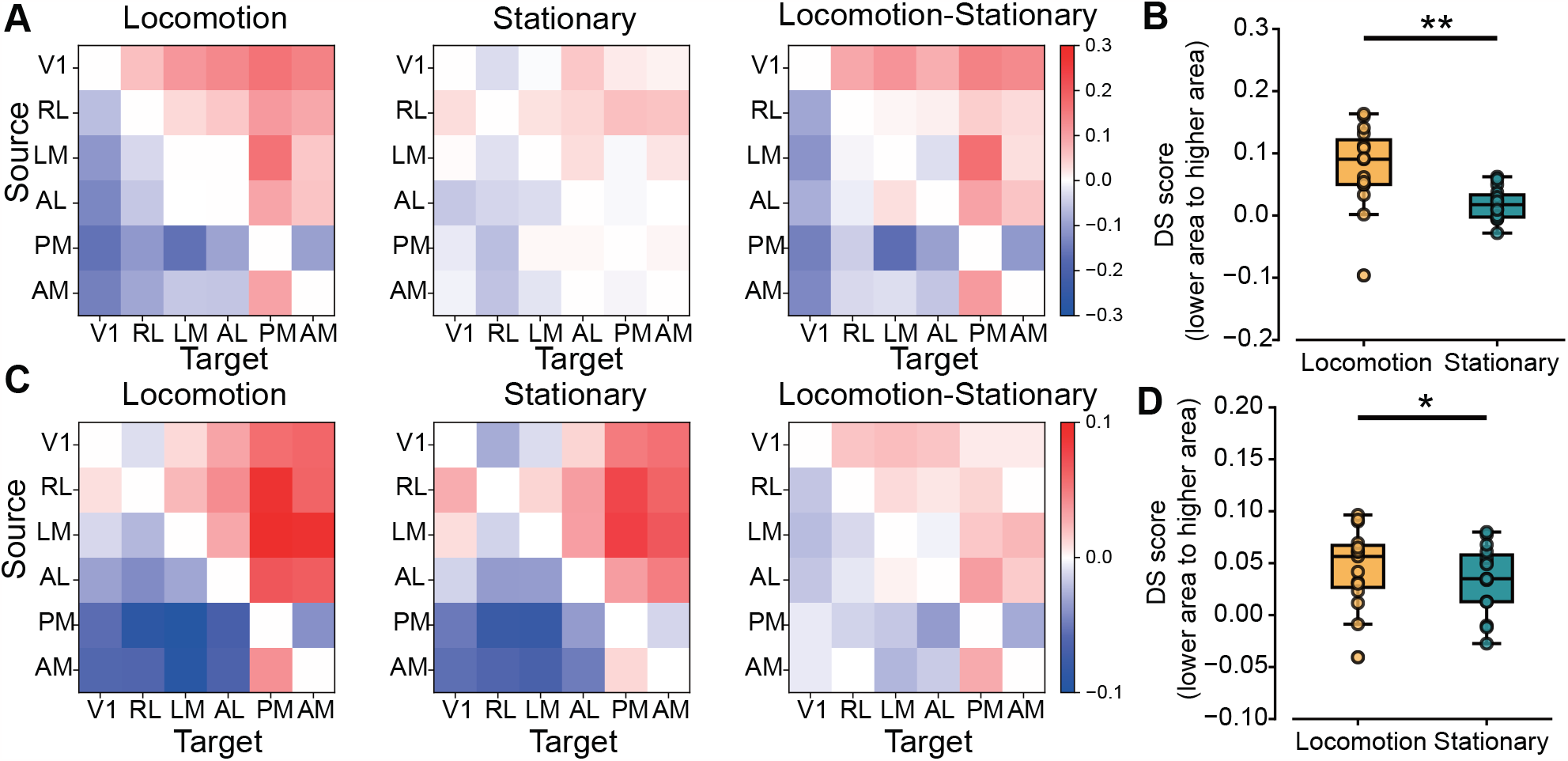
Feedforwardness quantified using alternative measures. **A**, Directionality score measured using reachability during locomotion and stationary states, and for the locomotion-minus-stationary difference. **B**, Directionality score from anatomically lower to higher areas during locomotion and stationary states. (Paired t-test, *n* = 15 pairs, *p* = 0.007). **C**, Same as in **A**, but with directionality score measured using canonical correlation analysis. **D**, Same as in **B**, but with directionality score measured using canonical correlation analysis. (Paired t-test *n* = 15 pairs, *p* = 0.013).

**Extended Data Fig. 13.**
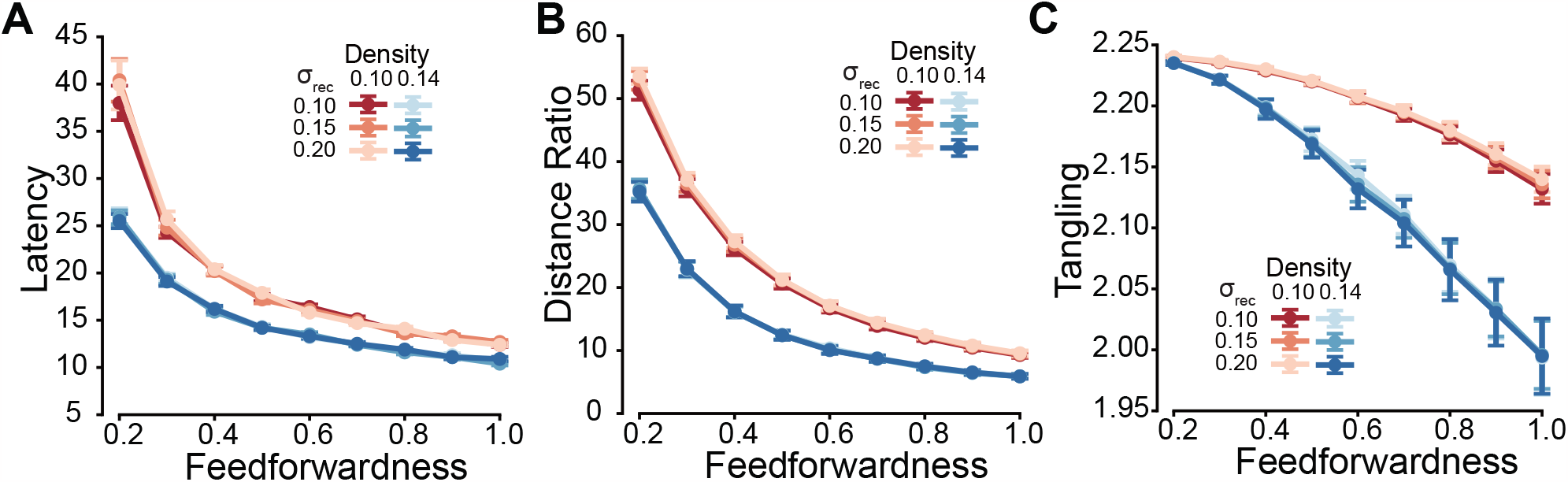
Feedforward architecture accelerates decoding and straightens neural dynamics. **A**, Decoding latency decreases as feedforwardness increases, under different densities and spatial scales of recurrent connectivity. **B**, Distance ratio decreases as feedforwardness increases, under different densities and spatial scales of recurrent connectivity. **C**, Tangling index decreases as feedforwardness increases, under different densities and spatial scales of recurrent connectivity. Points and error bars indicate mean ± SEM across 20 simulation repeats.

